# Epigenetic evolution of colorectal cancer and its microenvironment reveals new vulnerabilities

**DOI:** 10.64898/2026.08.10.743840

**Authors:** Alexandra Livanova, Luca Azzolin, Francesco Cambuli, Francesca Chemi, Chela James, Antonello Ferrazzano, Francesco Rusconi, Sabrina D’Agosto, Ilaria Mulas, Giovanni Randon, Michele Prisciandaro, Alessandra Raimondi, Filippo Ghelardi, Margherita Ambrosini, Paolo Manca, Federica Palermo, Monica Niger, Elisa Sottotetti, Antonia Martinetti, Marianna Maspero, Carlo Sposito, Paolo Ferrari, Luigi Antonio Lamparelli, Daniel Carrillo Bautista, Eugenia Ricciardelli, Fabio Simeoni, Luca Rotta, Clelia Peano, Vincenzo Mazzaferro, Trevor Graham, Filippo Pietrantonio, Andrea Sottoriva

## Abstract

Epigenetics is central to tumorigenesis, but the co-evolution of the cancer epigenome and its microenvironment is severely understudied. Here, we measure chromatin accessibility and transcriptome, at single cell resolution, of a set of normal colon, primary colorectal cancers and metastases, and identify recurrent epigenetic alterations in tumour cells. We also found that the normal epithelium adjacent to the cancer had recurrent epigenetic alterations associated with inflammatory programs that were partially shared with tumour cells. Distinct tumour-intrinsic transcription factor binding programs were associated with differential abundance of malignant stroma cell identities. We then leveraged matched patient-derived organoids, to assess the functional impact of the most recurrent epigenetic alterations on cancer cell viability, using CRISPR interference. We found a set of epigenetic-driven cancer dependencies, related to developmental reprogramming and cellular stress resilience, representing new potential therapeutic targets.

## INTRODUCTION

Epigenetic reprogramming is now recognized as a hallmark of cancer, acting alongside genetic alterations to drive malignant transformation and tumor evolution. The clinical relevance of epigenetic dysregulation is underscored by the FDA approval of several epigenetic drugs, including DNA methyltransferase inhibitors and histone deacetylase inhibitors^1,2^. These agents act as global chromatin modifiers with limited locus specificity and broad downstream effects. This lack of precision largely reflects the absence of an evidence-based catalogue of recurrent, functionally validated epigenetic driver events in cancer, in contrast to the well-defined genetic drivers that underpin most targeted therapies.

In colorectal cancer (CRC), recurrent genetic alterations (including those affecting *APC*, *TP53*, and *KRAS*, among the most common) confer niche independence and tolerance to genomic instability^3–7^. Historically, such mutations have been hard to target, and most CRC patients are routinely treated with broad spectrum chemotherapies.

At the transcriptional level, widespread de-differentiation is associated with the emergence of canonical (*e.g*., resembling intestinal stem cells (ISCs)) and non-canonical states including regenerative programs linked to fetal development^8,9^ and transiently observed during injury repair in healthy adulthood^10–12^. Such states display stereotypical spatial patterning across the tumour core – invasive margin axis since the earliest stages of disease^13,14^, and dependency on tumour-extrinsic signalling from the stroma^15–18^. Suppression of the anti-tumour immune microenvironment is further linked with progression to local invasion and distal metastasis^19–24^.

Somatic chromatin accessibility alterations (SCAAs) at regulatory elements *(e.g*., promoters and enhancers) are strongly associated with persistent gene expression programs during normal development and disease, and considered a key feature of long-term epigenetic memory^25–27^. Leveraging ATAC-seq (e.g., Assay for Transposable Accessible Chromatin^28^ in spatially-resolved sampling of individual multicellular clones (*e.g.*, glands) of CRC cells *vs.* normal mucosa, in earlier studies we found reduced accessibility at binding sites of interferon-regulated transcription factors (TFs) and of the chromatin insulator CTCF, as well as enhanced representation of developmental homeobox TF motifs^29^. Initial applications of single cell modalities (*e.g.*, scATAC-seq^30^) have begun to characterize the chromatin accessibility profile of individual CRC tumour and stromal cells^31,32^, but with limited representation of advanced stages of disease. More broadly, altered chromatin regions have been reported so far, but without functional validation of their biological relevance.

Here, we performed simultaneous single-nucleus ATAC- and RNA-seq (snATAC- and snRNA-seq using the 10x Genomics Multiome assay) on 33 primary CRCs, 11 CRC metastases, and 7 distant normal colon samples (including 7 patients with matched pairs of CRC specimens). We investigated the interplay between genetic and epigenetic alterations across CRC stages, from non-malignant tumour-adjacent epithelial cells to primary tumours and metastases. Transcription factor binding analyses identified distinct tumour-intrinsic regulatory programs displaying association with specific cell subpopulations from the malignant stroma, and reflecting microenvironmental variance across primary CRC and metastases. Finally, we performed functional CRISPR interference (CRISPRi) screens of a subset of promoters and enhancers, recurrently gaining chromatin accessibility in tumour cells, to assess their contribution to cancer cell viability. Together, our study defines a set of candidate epigenetic drivers—recurrent chromatin accessibility alterations that regulate gene expression and are required for cancer cell survival *in vitro*.

## RESULTS

### Convergent evolution of somatic chromatin accessibility alterations (SCAAs) at promoters and enhancers

Extensive DNA sequencing studies have elucidated the distribution of CRC genetic alterations^4,33–35^. In contrast, information on SCAAs has been lagging, especially because bulk assays, though informative, do not disentangle chromatin accessibility in cancer cells *vs.* other cell identities across the tumour microenvironment. Moreover, earlier efforts to apply single-cell assays have mostly overlooked advanced stages (*e.g.*, metastases) and matched specimens, with the latter particularly relevant to investigate epigenetic stability across space and time^31,32^.

We assembled a cohort of CRC tissue samples comprising primary tumours (n=33) and metastases (n=11) from 37 patients undergoing biopsy or resection at the *Istituto Nazionale dei Tumori (*Milan, Italy) (**Fig. 1a**; **Extended Data Fig. 1, Extended Data Table 1**). Within this cohort, seven sample pairs represented matched specimens, including pre- and post-treatment primary tumours (n=5 pairs), and synchronous collections at multiple sites (n=2 pairs). In addition, seven samples of distant normal colon tissue were included as non-malignant controls.

**Figure 1.**
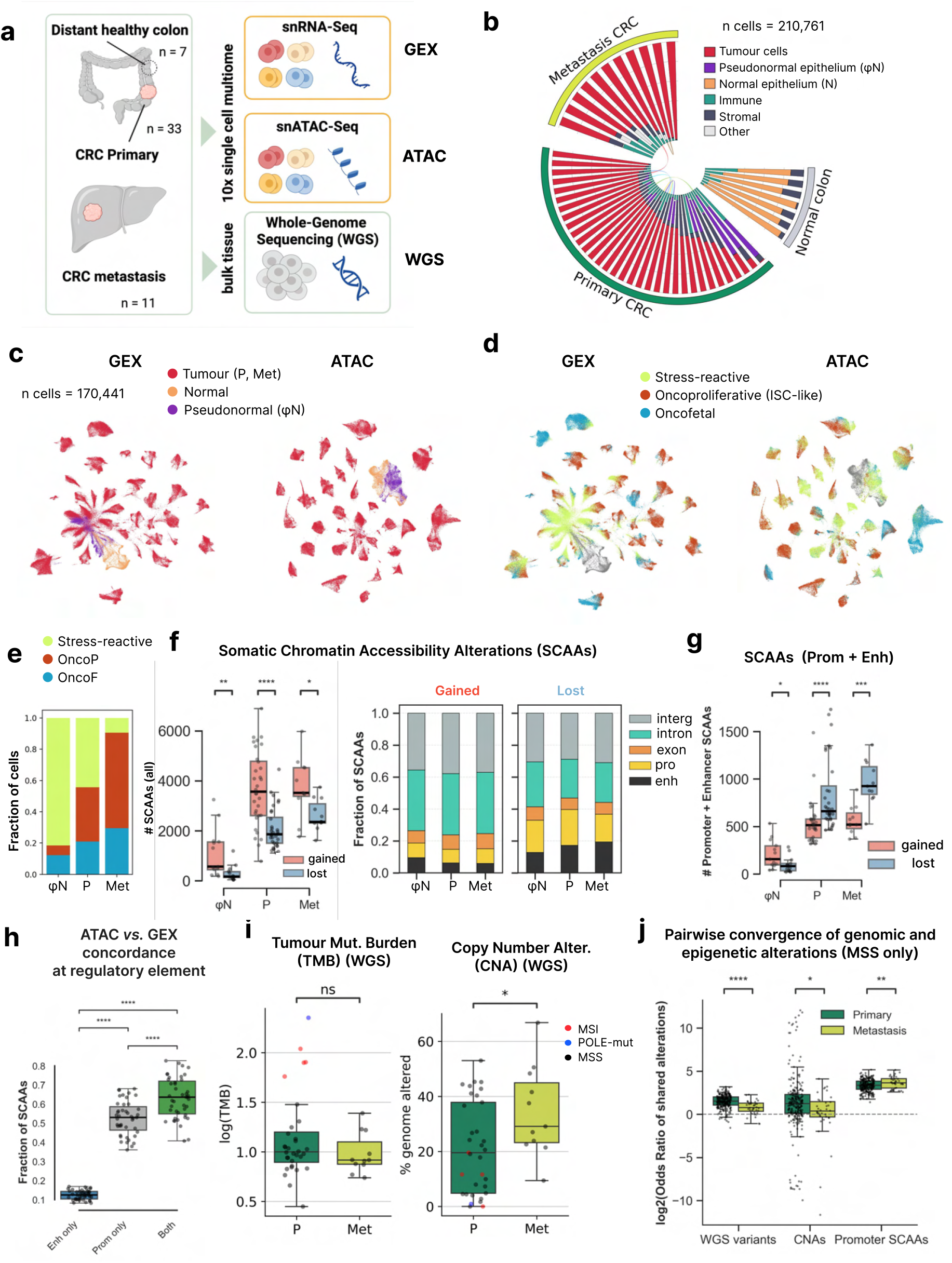
Convergent evolution of somatic chromatin accessibility alterations (SCAAs) at promoters and enhancers. **(a)** Experimental design. **(b)** Overview of the 10x multiome CRC cohort. Links connect matched samples from the same patient (See Extended Data Figure 1). **(c)** UMAP projections (GEX and ATAC) showing normal, pseudonormal and tumour epithelial cells. **(d)** UMAP projections showing major epithelial transcriptional states. **(e)** Abundance of transcriptional states across epithelial compartment **(f)** Number and genomic annotation of gained and lost somatic chromatin accessibility alterations (SCAAs).**(g)** Number of promoter- and enhancer-associated SCAAs. **(h)** Fraction of SCAAs exhibiting concordant changes in chromatin accessibility and expression of the associated target gene, shown separately for promoter-only, enhancer-only and jointly altered promoter–enhancer regulatory regions. **(i)** Tumour mutational burden (TMB) and fraction of genome altered (FGA) in primary and metastatic CRCs. **(j)** Paiirwise convergence of genomic and epigenomic alterations measured as odds ratios for shared single-nucleotide variants, copy-number alterations and promoter-associated SCAAs. Only MSS samples are shown, matched longitudinal samples removed.

Fresh-frozen tissues were profiled using the 10x Genomics Chromium Single Cell Multiome platform (see Methods), enabling simultaneous single-nucleus transcriptomic (gene expression, GEX) and chromatin accessibility (ATAC) profiling. In parallel, bulk whole-genome sequencing (WGS) was performed for each tumour sample to investigate the interplay between genetic and epigenetic changes in CRC (**Fig. 1a, Extended Data Fig. 2a**).

The single-cell multiome cohort comprised a total of 210,761 cells of which 147,596 were annotated as tumour (**Fig. 1b, c**; **Extended Data Fig. 2**), based on inferred single-cell CNV profiles closely recapitulating those obtained from matched bulk WGS (**Extended Data Fig. 3**); the remaining genomically-stable cells were assigned to specific cell identities according to established marker genes (**Fig. 1b, c, Extended Data Fig 2, Extended Data Table 2**). Additionally, 12 out of 33 primary CRC specimens included > 100 colon epithelial cells with unperturbed inferred CNV profiles (n= 14,364 in total), resembling those from normal distal specimens at the genomic level, but transcriptionally distinct (**Fig 1b, c; Extended Data Fig. 2, 3, 4**). We termed such cells ‘pseudonormal’ as, we reasoned, they were included in a subset of primary CRC specimens due to the nature of gross pathological sampling or dissection in tumours with irregular boundaries, in line with previous reports^36^.

Dimensionality reduction using UMAP embedding in either ATAC or GEX space revealed pronounced, and concordant, inter-patient separation among tumour cells, consistent with genome-wide patient-specific epigenetic/transcriptional profiles and copy number alterations (**Fig. 1c**). Tumour cells from matched samples derived from the same individual (n = 7 pairs), including pre- and post-treatment primary tumours or synchronous primary and metastatic lesions, clustered together, indicating a high degree of intra-patient similarity (**Extended Data Fig. 2b**). Furthermore, leveraging 38 gene sets from 12 previous studies on CRC transcriptional heterogeneity (**Extended Data Fig. 5a, Extended Data Table 2**), we classified pseudonormal and tumour cells into three main transcriptional clusters (see Methods). Based on manual annotation of the resulting hierarchical clusters, we termed such transcriptional states stress-reactive, reflecting an epithelial response to persistent inflammatory signalling (*e.g*., *RNF213^high^, HSP90AA1^high^, DUOX2^high^*)^37–39^, oncofetal (*e.g*., EMP1*^high^*, ANXA1*^high^, LAMC2^high^*) and oncoproliferative (*e.g.*, *LGR5^high^*, *RNF43^high^, MIKI67^high^*) (**Fig. 1d, Extended Data Fig. 5b)**. All three states were detected across stages, but the stress-reactive program was dominant in the pseudo-normal epithelium, whereas both oncofetal and oncoproliferative states were prevalent in CRC, jointly accounting for approximately 50% of the primary tumour cells, and more than 80% of metastatic (**Fig. 1e, Extended Data Fig 5c**).

Chromatin accessibility, as assessed by Tn5-mediated DNA tagmentation (ATAC)^28^, is unevenly distributed across the genome. Tn5 preferentially binds to nucleosome-free regions, in particular promoters and enhancers, which have an open chromatin conformation accessible to transcription factors (TFs). Single-cell ATAC can interrogate hundreds of individual cells from distinct lineages and, especially if combined with simultaneous 3’ scRNAseq, can disentangle cell-type specific profiles, which are obscured in bulk assays. Still, due to data sparsity, and near binary readout (*e.g.*, each allele is either tagged or not by Tn5), is impractical to measure single-cell accessibility variation at individual loci^30^. Thus, following best practices, we pseudo-bulked epithelial cells assigned to normal, pseudonormal, primary tumour and metastases, and generated a shared reference of all possible chromatin accessibility peaks within our study, restricting to those detected in at least two samples, for a total of approximately 62,000 putative peaks. We pooled the seven normal samples, grossly dissected distally from tumour sites, and whose global profile is highly overlapping upon dimensional reduction (**Fig. 1c)**, to generate a unified colon-specific control set (see Methods). Upon pseudobulking each sample, we run a differential accessibility test to nominate somatic chromatin accessibility alterations (SCAAs) in individual tumour or pseudonormal specimens against the pooled normal epithelial cells, as control **(Extended Data Fig. 2f-g**). In comparison to normal, both primary and metastatic CRC displayed a significantly higher number of gained *vs*. lost SCAAs at the genome-wide level (approx. 3,000 gained *vs.* 2,000 lost SCAAs on average), with pseudonormal (φN) cell fractions displaying a similar pattern but substantially lower absolute numbers of events (*e.g*., approx. 1,000 gained *vs.* 600 lost SCAAs) (**Fig. 1f**). In contrast, P and Met, but not φN cell fractions, presented a reversed ratio at gene regulatory elements, with tumour cells losing about 600-800 SCAAs at promoters and, even more markedly at enhancers, and gaining only approx. 400-500 as a mean value (**Fig. 1g, Extended Data Fig 6a-c**). Such quantifications on chromatin accessibility sites across CRC stages shed light on longstanding observations on altered nuclear architecture^43–45^ and, especially, on focal DNA hypermethylation at promoter CpG islands within the context of global hypomethylation^29,46–48^. We also split each tumour sample according to the main transcriptional states (stress-reactive, oncofetal and oncoproliferative) and found that SCAA distributions and pairwise similarities were more strongly associated with patient identity than with transcriptional state (**Extended Data Fig. 6d, e**), in line with a recent report on tumour-intrinsic genomic/phenotypic traits across spatially defined transcriptional regions in early CRC lesions^49^.

Nucleosome-free regions are necessary but not sufficient for TF binding, and productive engagement by RNA Pol II^50^. Yet, little is known on the relationship between chromatin accessibility and transcription in clinical cancers^29^. We assessed the degree of convergence between SCAAs and corresponding differentially expressed genes (DEGs), finding that, on average, about 50% of SCAAs at promoters are associated with concordant changes in mRNA levels, further rising to approx. 65% if SCAAs co-occur in matched promoters and enhancers (**Fig. 1h**) (as found in 30% of promoter SCAAs as mean value) (**Extended Data Fig. 6f**).

A small set of mutations and copy number alterations (CNA) are recurrently found in CRC patients and known to alter functionally validated cancer driver genes (*e.g*., APC^mut^, TP53^mut^, KRAS^mut^, 20q^gain^, 8q^gain^, 18q^loss^)^4,33–35^. Here, we quantified the odds ratio of shared alterations across random pairs of tumours, considering protein-affecting somatic variants, CNAs, and promoter SCAAs, as measured by WGS or scATAC-seq (**Fig. 1i**). Promoter SCAAs exhibited a two- to three-fold higher odds ratio of sharing than either mutations or CNAs, demonstrating that these epigenetic alterations are substantially more recurrent across tumours. Notably, metastases showed an even higher odds ratio of shared promoter SCAAs than primary tumours, indicating reduced epigenetic heterogeneity and supporting positive selection of specific SCAAs during CRC progression (**Fig. 1j, Extended Data Fig. 6g**).

Thus, across CRC stages we found a global increase in chromatin accessibility, but a net loss at promoters and enhancers, with genomically stable pseudonormal cells adjacent to tumour displaying a more modest increase, yet uniformly across genomic regions. On average, about half of the SCAAs occurring at promoters were associated with concordant changes in gene expression, approaching two thirds in the presence of matched enhancer SCAAs. Finally, we observed substantial recurrence of promoter SCAAs across primary CRCs and metastases, consistent with convergent evolution during CRC progression.

### Frequency and stability of somatic chromatin accessibility alterations (SCAAs) across CRC stages

In contrast to the well-annotated catalogue of genetic alterations^4,33–35^, no systematic frequency analysis of chromatin altered sites across CRC patients is available. Thus, we sought to identify recurrent chromatin accessibility alterations (*e.g.*, either gained or lost SCAAs) across our CRC cohort. Restricting the analysis to 36 of 37 patients with more than 100 tumour cells profiled by scATAC-seq, we quantified the recurrence of promoter SCAAs and somatic nucleotide variants (SNVs) by grouping genes according to the number of patients in which they were altered, using 4-patient intervals (**Fig. 2a; Extended Data Fig. 7a**). As expected for CRC, we found only 16 genes mutated in at least 25% of patients (*e.g.*., >8 out 36), including key driver genes like *APC* and *TP53* (>70%) and *KRAS* (>25%). In contrast, the number of genes affected by SCAAs in more than a quarter of patients was approx. 12-fold higher (*e.g*., 399 genes with promoter SCAAs in >8 out of 36 pts.). Moreover, the frequency of SNVs and promoter SCAAs appears to be inversely related (*e.g*., if a gene is frequently mutated is unlikely to be affected by promoter SCAAs and *vice versa).* Focusing on SCAAs affecting promoters or enhancers with concordant changes in mRNA expression for the corresponding genes, and occurring in at least 25% of our cohort (*e.g.,* >8 out 36 patients), we found 104 genes with gained SCAAs, and 258 genes with lost SCAAs, that have not been previously implicated in tumorigenesis through genetic mutations (**Fig. 2b** displays the 25 most recurrent loci with either gained or lost SCAA localized at promoters and/or enhancers; **Extended Data Fig. 7b** is an analogous visualization restricted to known genetic drivers^34^, **Extended Data Table 3**). Gene set enrichment analysis found that highly recurrent genes affected by gained SCAAs at regulatory regions, are over-represented in pathways related to unsaturated and long-chain fatty acid metabolism, as well as epithelial development and morphogenesis, while lost SCAAs are associated with metal binding and transport across cell membranes (**Fig. 2c, Extended Data Table 4**). Collectively, such pathway-level associations point to loss of genes expressed in mature colonocytes, mediating ions and metal trafficking across the colonic lumen (*e.g*., *SCNNB1*, *SLC4A4*, *MT1E*, *MT1G*, *MT1M*)^39,51–54^ and upregulation of developmental transcription factors (*e.g*., *MSX2*, *PITX2*, *PDX1*)^54–57^ and genes responsible for cell membrane remodelling, cell adhesion, and extracellular matrix interactions (*e.g*. *TGFBI*, *PHLDA1*, *ITGA2*, *CLDN1*, *CDH3*)^58–62^. Although the size of our current cohort is suboptimal for performing robust comparisons across stages, we report several SCAAs with differential recurrence between primary tumours and metastases (**Fig. 2d; Extended Data Fig.7c**). Promoter SCAAs more prevalent in metastasis samples (>30% vs. primary CRC) were preferentially enriched in genes previously implicated in metastatic progression, including *MSX2*, *PITX2*, *TBX3*, *ACOT1*^63–66^. Conversely, genes associated with normal epithelial identity and function, such as *ALPI*, *ALPL*, *ADH1C*, and *MUC3C*^67,68^, or *DLC1* and *ARHGAP20* tumour suppressors^69,70^ were more often affected by promoter loss in metastases compared with primary tumours. This pattern is consistent with enhanced de-differentiation and increased cellular plasticity in metastatic lesions.

**Figure 2.**
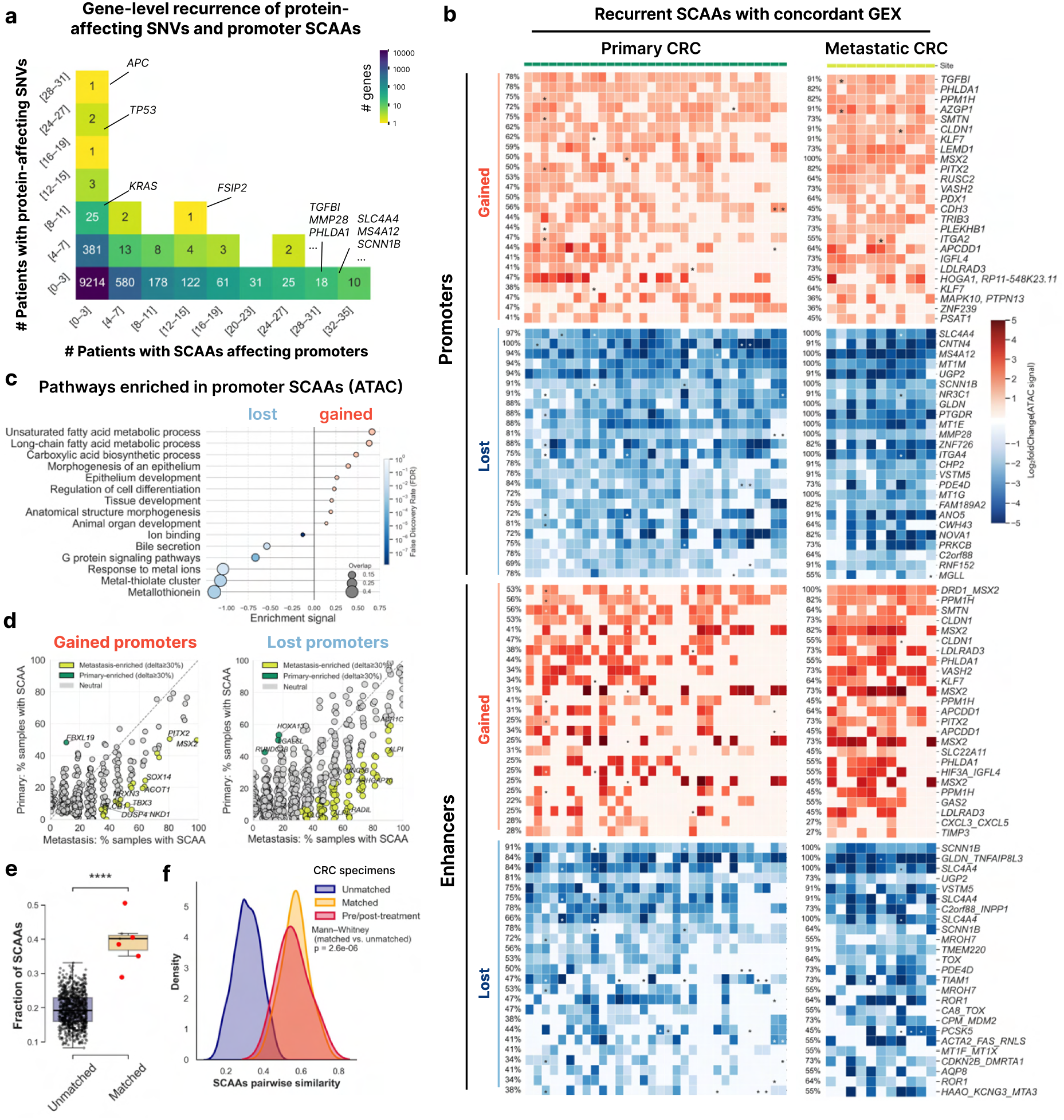
Tumor-specific SCAAs are highly recurrent, temporally stable epigenetic alterations with distinct patterns in primary and metastatic tumors. **(a)** Relationship between genetic (somatic variants) and epigenetic (SCAAs) regulation across cancer driver genes **(b)** Top recurrent SCAAs (|log₂ fold change| > 1, padj < 0.01) affecting promoters and enhancers relative to normal colon epithelial cells. Shown are SCAAs recurrent in at least three patients. Enhancers with a corresponding affected promoter are indicated. Asterisks denote the presence of somatic variants in the associated gene. Same heatmap for known cancer-related genes is presented in Extended Data Figure 7b. **(c)** Pathway enrichment in genes recurrently affected by SCAAs **(d)** Recurrence of promoter-associated SCAAs in primary and metastatic tumours. **(e)** Fraction of shared SCAAs in matched *vs*. unmatched CRC samples. Red dots stand for pairs of pre-/post treatment samples from the same patient. **(f)** Pairwise similarity of epigenetic profiles between matched vs. unmatched CRC samples, measured as Spearman correlation of log2 fold change values for each SCAA

Epigenetic stability appears to be context-dependent, ranging from developmentally established events, faithfully inherited across cell divisions throughout the life span, to fully reversible short-term perturbations induced by acute infections or inflammation^26,71–73^. To evaluate SCAAs temporal stability during CRC evolution, we exploited our set of seven pairs of matched samples collected from the same patients, including five paired pre- and post-treatment primary tumours, one treated pair of liver metastases from distinct lobes, and one synchronous untreated pair of primary and metastatic CRC. Matched samples from the same patient shared, on average, 37% of SCAAs, compared to 19% shared SCAAs in random unmatched pairs (**Fig. 2e; Extended Data Fig. 7d, e**). Furthermore, correlations of log_2_ fold changes of SCAAs were significantly higher in matched pairs than in random unmatched pairs, indicating that the overall chromatin accessibility profile remains relatively stable over time within individual patients (**Fig. 2e, f; Extended Data Fig. 7d, e**).

Furthermore, to explore the relationship between genomic and epigenomic intratumoral heterogeneity, we focused on sample 002_S1_T, which contained two clearly distinct CNA-defined tumour subclones identified by single-cell CNA inference (**Extended Data Fig. 8a–c**). Tumour cells assigned to each CNA cluster were pseudobulked separately, and chromatin accessibility profiles were normalized using clone-specific copy-number estimates prior to SCAA calling against pooled normal epithelial cells. Despite detected copy-number differences in specific segments, chromatin accessibility alterations were highly conserved between subclones: 70.9% and 46.0% of SCAAs identified in C1 and C2, respectively, were shared, and accessibility effect sizes showed strong concordance across clusters (Spearman ρ = 0.72, *P* < 0.001; **Extended Data Fig. 8d, e**).

Shared SCAAs were modestly enriched within genomic regions exhibiting conserved copy-number states (Fisher’s exact test, OR = 1.19), indicating that common CNA events contribute to, but do not fully explain, the stability of chromatin accessibility alterations across tumour subclones (**Extended Data Fig. 8f, g)**.

Hence, genetic variants and chromatin alterations affect orthogonal gene sets. We have compiled an atlas of regulatory elements with recurrent alterations to chromatin accessibility across CRC patients and stages. Chromatin accessibility losses occur at genes involved in intestinal differentiation and transport across the colonic lumen, while gained SCAAs target genes associated with developmentally related alterations to membrane remodelling, cell adhesion, and extracellular matrix interactions. A subset of such events appears at increased frequency in met vs. primary CRC. Finally, we found that chromatin alterations retain substantial temporal stability during CRC evolution.

### Analysis of tumour-adjacent pseudonormal cells reveals inflammation-driven chromatin alterations shared with CRC

Disentangling the contribution of tumour intrinsic and extrinsic oncogenic pathways in the CRC clinical scenario is complicated by their spatiotemporal overlap. Having previously observed a fraction of epithelial cells with unperturbed inferred CNV profile within 13 out of 33 primary CRC specimens (n= 10,405 in total) (**Fig 1b,c; Extended Data Fig. 2, 3, 4**), we reasoned that such pseudonormal (φN) cells experience extrinsic oncogenic signalling, due to their spatial proximity, but are unaffected by genetically driven intrinsic cancerogenic mechanisms. In contrast to tumour cells, experiencing a global increase in chromatin accessibility but a reduction at regulatory elements, φN cells display a moderate increment throughout the genome, including at promoters and enhancers (**Fig. 1f,g**). Our initial observations at the whole cohort level were confirmed by UMAP visualization of three matched pairs of primary CRC and anatomically distant colon specimens, including all three cell populations: normal (N), primary (P) CRC and adjacent pseudonormal (φN) (**Fig. 3a**). φN cells approximate normal cells in ATAC modality, but they cluster in proximity to P cells in gene expression (GEX), suggesting a greater shift in transcriptional *vs*. chromatin accessibility programs. Upon pseudobulking φN cells belonging to individual specimens, we performed differential chromatin accessibility analysis against epithelial cells from distant normal colon tissue and searched for shared SCAAs between φN and adjacent P cells. We found substantial global concordance (*e.g*., approx. 30% of gained and 60% of lost SCAAs in φN are shared with P), which was reduced to 10-15% *circa* at promoters and enhancer (*e.g.*, approx. 10% of gained and 15% of lost SCAAs at regulatory elements in φN are shared with P) (**Fig. 3b**). We annotated the resulting SCAAs and focused on promoter- and enhancer-associated regions that were gained or lost in φN cells. Across 12 patients, we identified 19 recurrently gained and 7 recurrently lost promoters present in at least three samples (*e.g*., 25% our cohort) (**Fig. 3c**; a black dot indicates a shared SCAA *vs.* matched primary CRC). Several promoters, recurrently gaining accessibility in φN cells, are known to control the expression of important mediators of colonic inflammation, including *MAPK10, OLFM4, DUOX2, IL1RN, PTGS2, MAP3K10,* and *ANGPTL4*^37–39^, while in the smaller promoter set loosing accessibility, there are key colonic transporters for water and ions, like *AQP8* and *MT1G,* whose function is affected by colitis^39,51,74,75^. Based on pathway enrichment analysis of genes linked to promoter SCAAs observed in at least two pseudonormal samples (**Fig 3d, Extended Data Table 4**)), recurrently gained SCAAs were enriched for multiple inflammatory and immune-related pathways, including eicosanoid metabolism, biosynthesis of specialized pro-resolving mediators (SPMs), IL-4/IL-13 signalling, and interleukin signalling. In contrast, recurrently lost SCAAs exhibited a more restricted enrichment profile, with the strongest enrichment observed for metallothionein- and metal ion–associated pathways.

**Figure 3.**
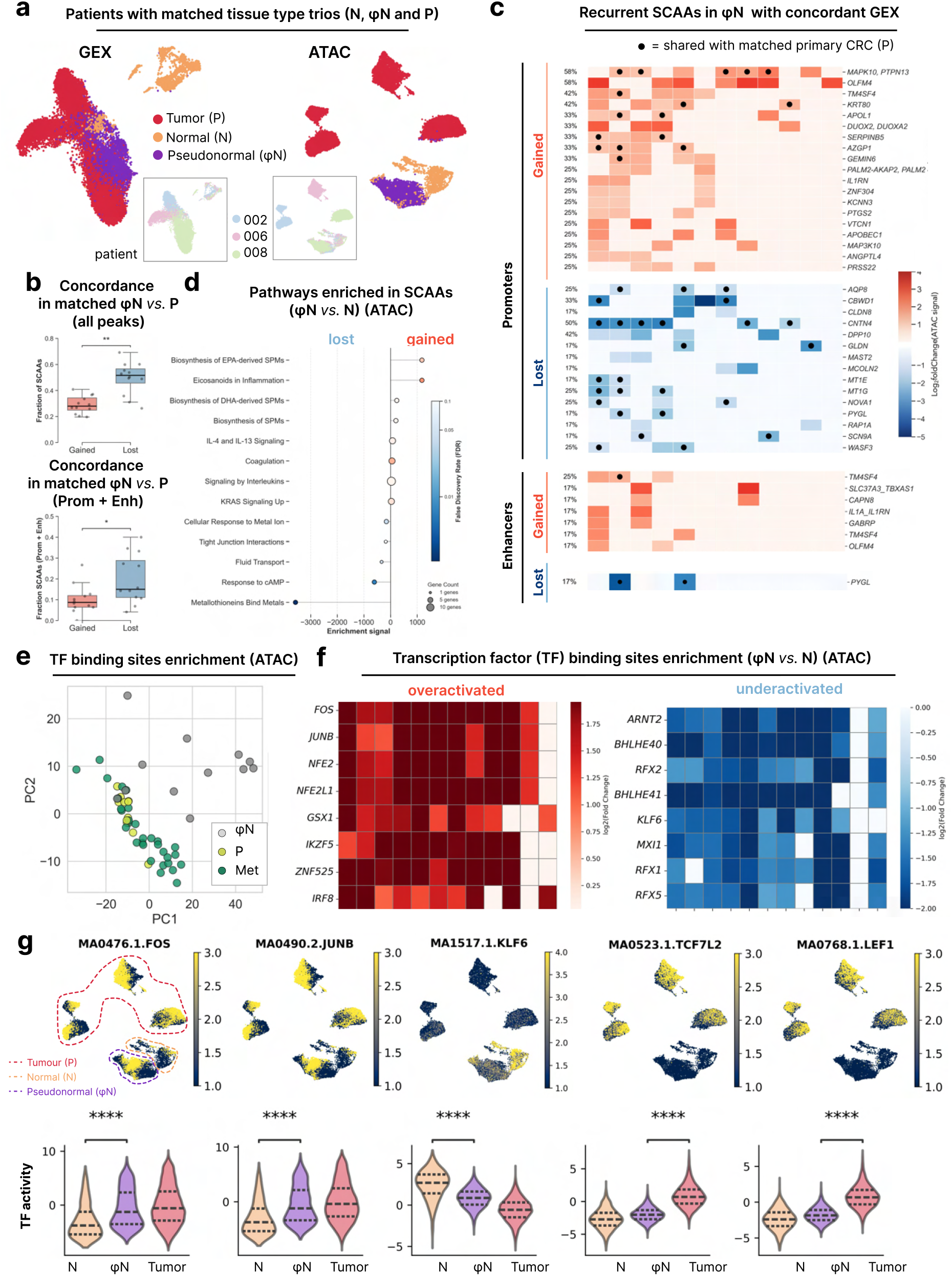
Epigenetic and genomic alterations in pseudonormal epithelial cells adjacent to colorectal cancer (CRC). **(a)** UMAP projections (GEX and ATAC) showing normal, pseudonormal and tumour epithelial cells from three patients (002_S1, 006_S1, 008_S1) **(b)** Fraction of SCAAs inferred in pseudonormal epithelium concordant with SCAAs from matched primary tumour, all SCAAs (*top*) and promoter+enhancer regions (*bottom*) **(c)** Top recurrent SCAAs (|log₂ fold change| > 1, padj < 0.01) affecting promoters and enhancers in pseudonormal epithelial cells compared with distant normal epithelial cells. SCAAs were considered recurrent if affecting promoters in ≥3 patients or enhancers in ≥2 patients; enhancers with matched affected promoters are shown. Marker indicates presence of the corresponding SCAA in tumour cells from the matched sample**. (d)** Pathway enrichment of genes with promoters affected by SCAAs in pseudonormal epithelium relative to distant normal cells.**(e)** Principal component analysis (PCA) of transcription factor enrichment scores, showing pseudonormal epithelium, primary and metastatic tumour cells. **(f)** Transcription factor (TF) motif enrichment in SCAAs gained and lost in pseudonormal epithelium (TFs recurrent in ≥10 samples are shown). **(g)** Top, UMAP projection of single-cell chromatin accessibility profiles for a subset of patients (002_S1, 006_S1, 008_S1) colored by motif deviation (chromVAR deviation score calculated with pychromVAR) for representative transcription factor motifs (*FOS, JUNB, KLF6, TCF7L2*, and *LEF1*). Bottom, distribution of motif deviation scores across cells from each tissue type (N, φN, and tumour). Horizontal dashed lines indicate the median and quartiles.

To gain insights into coordinated signalling and transcriptional programs across CRC stages^76,77^, we performed transcription factor motif enrichment analysis using known binding motifs for 1,165 transcription factors from the CIS-BP database in SCAAs across φN, primary or met CRC cells (log₂ fold change > 1, adjusted P < 0.01), using peaks enriched in distant normal epithelial cells as background (log₂ fold change < −1, adjusted P < 0.01). From this analysis, we identified 767 transcription factors showing significantly altered motif activity in at least five samples across pseudobulked φN cells adjacent to CRC, primary tumours, and metastatic lesions. Principal component analysis (PCA) based on the first two components of the transcription factor activity matrix revealed a clear separation between pseudonormal and tumour cell samples, as well as greater inter-individual variance across φN cell populations, as expected for cells retaining a strong epigenetic memory of their somatic identity (**Fig. 3e**). We initially focused on transcription factor binding motifs (TFBS) enriched in pseudonormal cells adjacent to CRC *vs*. normal cells from anatomically distinct colon specimens. In almost all patients, φN cells exhibited overrepresentation of AP-1 sites, putatively bound by *FOS*, *JUNB, NFE* and *NFE2L1*, and depletion of E-box-like motifs associated with bHLH-related TFs, like *ARNT2*, *BHLHE40*, *BHLHE41*, and *MXI1*, as well as underrepresentation of RFX-family sites (*RFX1/2/5*) and *KLF6*-related motifs (**Fig. 3f, Extended Data Table 5).** Among the TFs with inferred increased accessibility, *FOS* and *JUNB* are canonical members of the *AP-1* complex which is a primary mediator of stress-activated early-response transcriptional programs, with a major role in chronic colonic inflammation (*e.g*., ulcerative colitis)^38,73,78^. In the opposite direction, TFs predicted to lose accessibility are known to promote cell cycle exit (*KLF6*, *MXI1*)^79–81^, regulate circadian rhythms (*BHLHE40/41*)^82,83^, whose disruption facilitates oncogenic transformation in intestinal organoid models^84^, and transcription of MHC class I molecules and β2-microglobulin (RFX5)^85^.

By visually inspecting and quantifying the enrichment for individual TF binding sites at single-cell level, across three matched pairs of primary CRC and anatomically distant colon specimens (**Fig. 3a**), we confirmed a large fraction of φN cells displaying a significant enrichment for AP-1 (*FOS/JUNB*) sites, and *vice versa*, a depletion of *KLF6* motifs, in comparison to distal normal cells (**Fig. 3g**). We also observed similar representation of *FOS*, *JUNB*, and *KLF6* target sequences between φN and primary CRC cells, indicating that such chromatin alterations are predominantly driven by extrinsic signalling rather than intrinsic genetic alterations. In contrast, TCF/LEF motifs (*TCF7L2*, *LEF1*) are markedly overrepresented in tumour cells, while pseudonormal cells retain basal levels, supporting mutagenesis of WNT family genes (*e.g*., *APC*^mut^) as main cell-autonomous activation modality.

Thus, tumour-adjacent epithelia may be oversusceptible to transformation due to extrinsic inflammatory signalling, providing an epigenetic base for the field effect hypothesis^86,87^ in CRC, complementing previous studies on DNA methylation in bulk cellular samples^88–90^.

### The canonical CRC transcription factor binding program displays a near-binary modulation related to translational-stress

We extended our exploration of transcription factor binding programs in tumours by focusing on primary and met CRC samples. To gain a deeper understanding, we further restricted the analysis to 361 TFs, exhibiting significantly altered motif enrichment in >25% of our patient cohort (>9 out 36 samples) (**Fig. 4a** and **Extended Data Fig. 9a**), and performed hierarchical clustering (**Extended Data Fig. 9b, c)**. We obtained four main TF subsets which were manually annotated based on the most prevalent motifs, and the known biological functions of the related transcription factors as follows: CRC core (*TCF*/*LEF*, *AP-1*) featuring a small group of WNT- and inflammation-driven TFs (*e.g*., *LEF1, TCF7L2, FOS, JUNB*)^78,91^; Fetal development (*HOX, FOX, SOX*) encompassing a broad selection of master regulators of gastrointestinal progenitors (*SOX17, FOXA2, HNF4A, HNF1B, PDX1, HHEX, ONECUT1, SATB2, GFI1, NEUROD1* and *SOX2*)^92–94^; Translational stress (*ATF, CREB, TFEB*) spanning a variety of key regulators of protein synthesis and autophagy (*ATF4, ATF6, ATF6B, XBP1, CREB3, CREB3L1, TFEB* and *MITF*)^95,96^; Colon differentiation (*KLF, SP, ZBTB, ZNF*), including TF promoting intestinal differentiation (*HES1, KLF4, KLF5, KLF6, ZBTB33, NR3C2*)^97^, as well as epigenetic factors enforcing lineage stability (*CTCF, DNMT1, MBD2*)^98,99^ (**Fig. 4a**, **Extended Data Table 5**). Upon dimensionality reduction, we noticed two main groups of specimens, with one of them enriched in metastases (**Fig. 4b**). The CRC core (*TCF*/*LEF*, *AP-1*) was the only main TF subset to retain an obvious motif enrichment in both groups of samples, in comparison to normal cells, although with a significant shift between the two cohorts. In contrast, only one of the specimen groups featured overrepresentation of Fetal dev TFs (*HOX/FOX/SOX*), as well as underrepresentation of Colon Diff TFs (K*LF, SP, ZBTB, ZNF*), with the other group retaining such motif profiles close to basal levels, but derepressing Translational stress TF motifs (*ATF, CREB, TFEB*). As our observations are consistent with a canonical CRC TF binding program, partially restrained by translational stress response, we termed the former ‘Hypercanonical’ and the latter ‘Hypocanonical’ (**Fig. 4c**).

**Figure 4.**
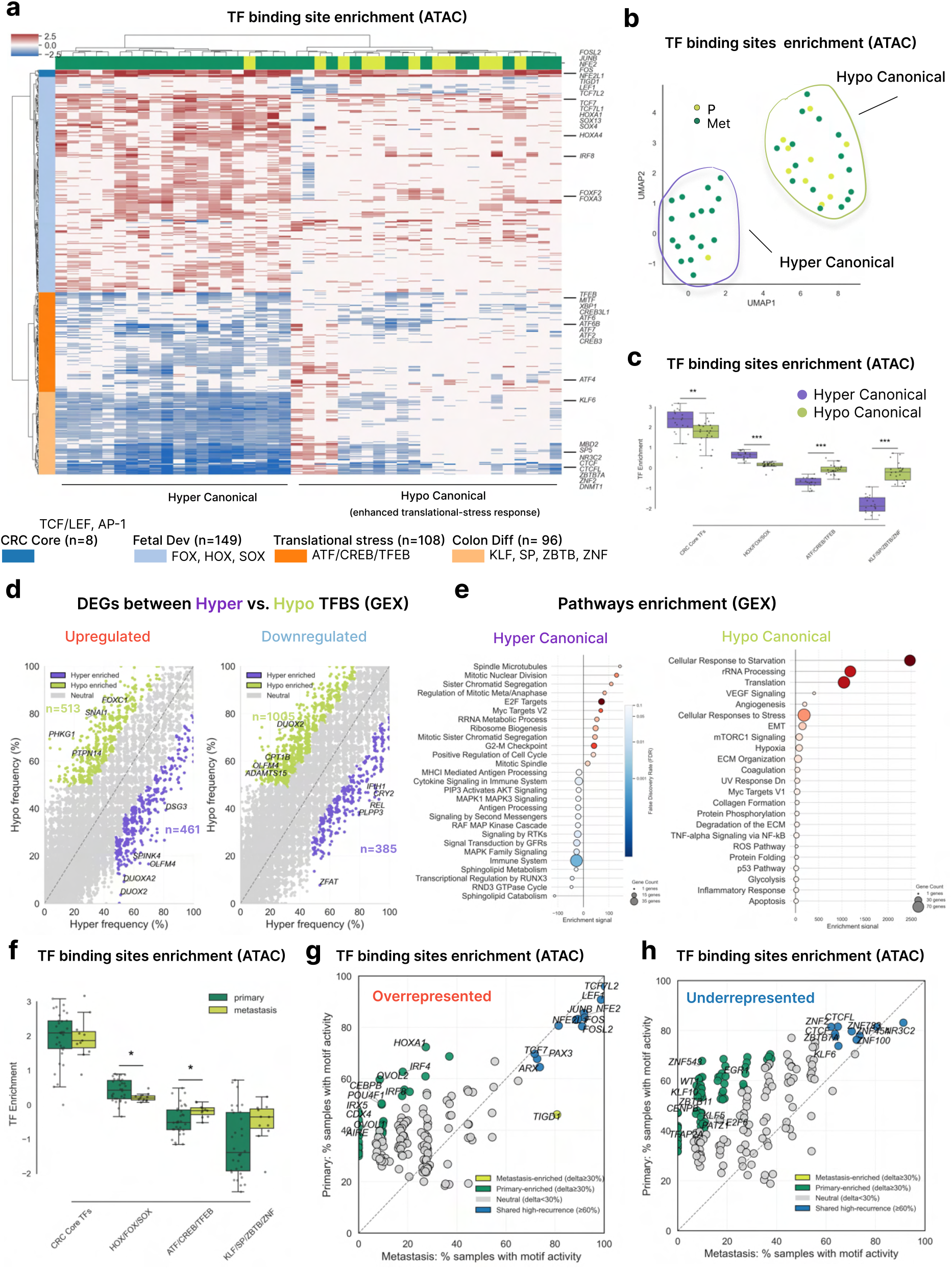
The canonical CRC transcription factor binding program displays a near-binary modulation related to translational-stress. **(a)** Motif enrichment analysis of binding sites corresponding to 361 TFs enriched in gained and lost SCAAs. TFs significantly upregulated or downregulated in at least nine patients are shown. TFs (rows) were ordered according to four clusters identified by hierarchical clustering (see also Methods; Extended Data Fig. 9). **(b)** UMAP projection of CRC samples based on TF cluster scores. The analysis reveals two distinct groups of samples, Hypo- and Hyper-Canonical. **(c)** Averaged TF enrichment scores for the four clusters in Hypo and Hyper-canonical groups. **(d)** Recurrent differentially expressed genes in Hypocanonical vs. Hypercanonical groups **(e)** Pathway enrichment of DEGs recurrent in Hypo- and Hyper-canonical groups of samples **(f)** Averaged TF enrichment scores for the four clusters, comparing primary and metastatic tumour samples.**(g)** Recurrence of the upregulated and **(h)** downregulated TFs in primary and metastasis tumours

We expanded our analysis by comparing DEGs frequency across Hyper- vs. Hypocanonical tumours, identifying 461 genes recurrently upregulated in the former and 513 in the latter, while 385 and 1005 were downregulated, respectively (**Fig. 4d, Extended Data Table 6**). Gene set enrichment analysis on recurrent DEGs between the two groups yielded an enrichment for pathways related to cell cycle progression and biosynthesis in Hypercanonical tumours. In contrast, Hypocanonical displayed an overrepresentation of genes associated with cellular response to starvation and angiogenesis (**Fig. 4e, Extended Data Table 4**).

The distinction between Hypercanonical and Hypocanonical tumours was not purely driven by the clustering of most metastatic samples in the latter, as the representation of the main TFBS clusters is only modestly altered in primary *vs.* met CRC, due to the broad variance displayed by primary tumours (**Fig. 4f**). Instead, a subset of primary tumours is characterized by TFBS profiles reminiscent of those observed in metastatic samples. Nevertheless, when we tested the frequency of TFBS enrichment between primary *vs.* metastatic specimens (**Fig 4g,h**), some were preferentially associated with the former, while only one binding site was preferentially represented in metastases, putatively bound by a transposable element derived gene, *TIGD1*, poorly characterized but recently reported as overexpressed in advanced colorectal cancer stages^97^. In summary, we observed two distinct transcription factor binding programs across colorectal cancers, termed Hypercanonical and Hypocanonical. The first state is associated with cycling and biosynthetically active tumour cells, featuring maximal activity of WNT and inflammatory TFs, as well as reactivation of developmental programs and repression of intestinal differentiation. Such state, more prevalent in primary CRC, is consistent with a previous chromatin mini-bulk profiling carried out by our group on micro-dissected individual tumour glands from an unrelated CRC cohort^29^. The second state is chiefly characterized by de-repression of translational-stress pathways, typical of a response to a microenvironment depleted in resources, like nutrients, growth factors or oxygen, as commonly experienced in metastatic settings but also in a subset of primary tumours, upon tissue damage by endogenous immunity or exogenous therapies.

### Dynamic epigenetic remodelling of the tumour microenvironment during CRC progression

As single-nucleus dissociation protocols from clinical fresh frozen specimens are not neutral to cell lineages^98,99^, and epithelial cells were our primary focus, we recovered a relatively lower fraction of cells from the tumour microenvironment (**Fig. 1b, Extended Data Fig. 2**). Coupled with the substantial diversity of stromal and immune cell populations, the limited number of cells per cell type precluded robust sample-level epigenetic analyses. Therefore, unlike normal epithelial, pseudonormal, and CRC tumour cells (**Fig. 1**), stromal and immune cell populations were analysed after aggregation across samples to characterize global cell type-specific chromatin accessibility programs.

Nevertheless, we recovered and systematically characterize 40,320 TME cells, including 13,903 stromal (**Fig. 5a**; crypt fibroblasts, CAFs, endothelial, lymphatic, pericytes, and glia) and 12,021 immune cells (**Fig. 5b**; T/B cells, plasma, myeloid and dendritic cells), as well as epithelial and other cell identities from distinct organs targeted by metastases. To define the regulatory programs underlying stromal and immune cell identities, we leveraged the ATAC modality to identify cell type-specific accessible chromatin regions. Peaks were aggregated within each cell population across all patients, marker peaks were identified by differential accessibility analysis, and transcription factor binding site (TFBS) enrichment was subsequently performed on cell type-specific regulatory elements (**Fig. 5c, d**). The resulting TF activity profiles recapitulated known lineage-defining regulators, including enrichment of *FLI1*, *ERG*, and *EBF1* in endothelial/perivascular compartments^100–102^; *TWIST1*, *CEBP* family factors and FOX proteins in fibroblast populations^103–105^; and *AP-1* complex members (FOS, JUNB) in CAFs (**Fig. 5c**). In immune cells, TFBS enrichment recovered canonical regulators of lymphoid and myeloid differentiation, including *TCF7L2*, *LEF1*, *RORA*, and *EOMES* in T-cell populations, *IRF4*, *IRF5*, and *IRF8* in B-cell and plasma-cell compartments, and *NFKB1/2, RELA, CEBPA*, and *CEBPB* in myeloid cells (**Fig. 5d**).

**Figure 5.**
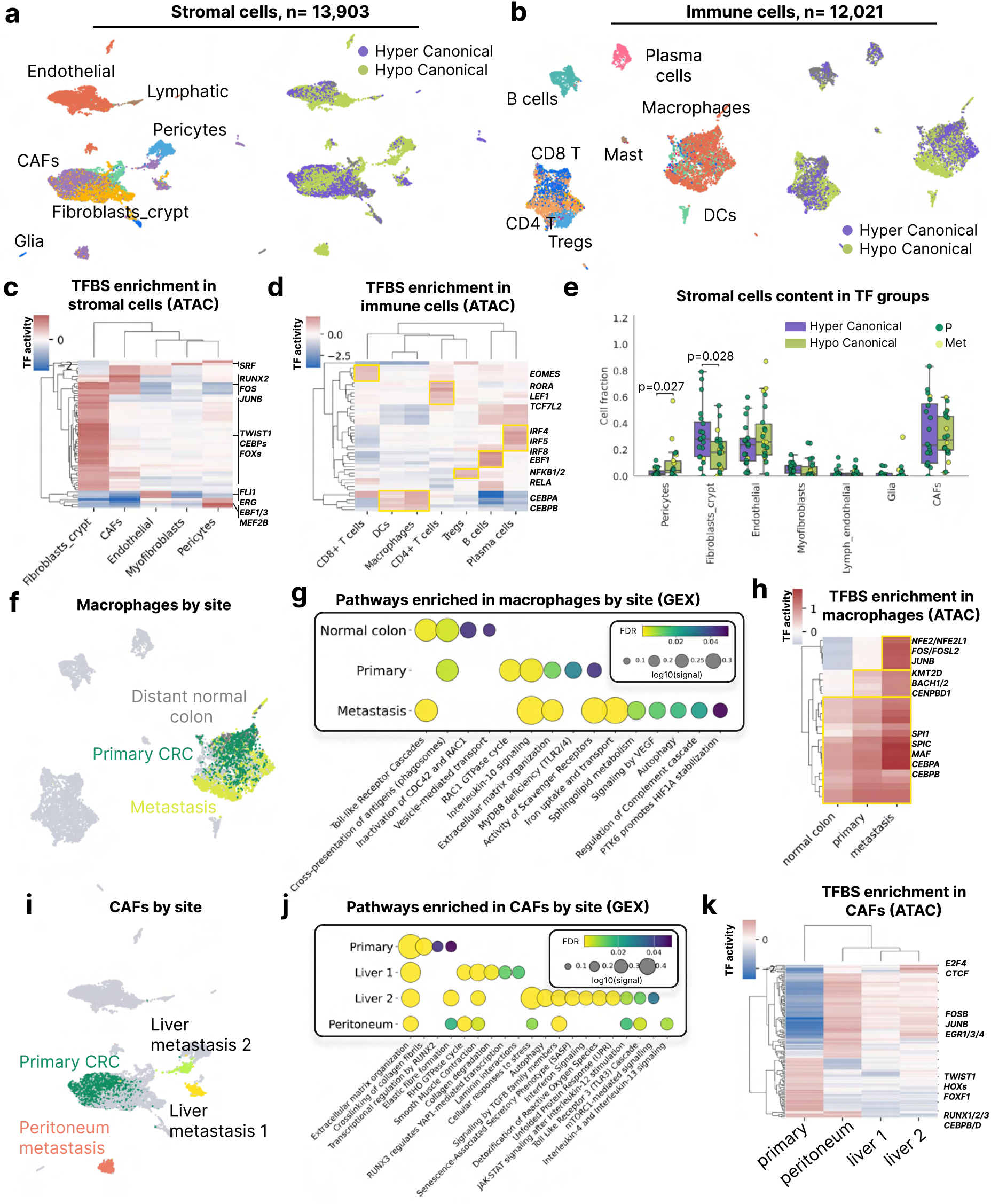
Dynamic epigenetic remodelling of the tumour microenvironment during CRC progression. **(a)** UMAP projection of scATAC-seq data showing 13,903 stromal cells across the CRC cohort. **(b)** UMAP projection of scATAC-seq data showing 12,021 immune cells across the CRC cohort.**(c)** TFs differentially enriched in marker peaks associated with stromal and **(d)** immune cell populations. For each cell type, motif enrichment was performed using marker peaks with all remaining peaks as background. **(e)** Stromal cells abundance in Hypo- and Hyper-canonical groups of tumour samples **(f)** UMAP projection of scATAC-seq data showing macrophages annotated by tissue site. **(g)** Pathway enrichment analysis of marker genes in macrophages from normal colon, primary CRC, and metastatic CRC. **(h)** TFs differentially enriched in marker peaks associated with macrophages from normal colon, primary CRC, and metastatic CRC. For each cell type, motif enrichment was performed using marker peaks with all remaining peaks as background. **(i)** UMAP projection of scATAC-seq data showing CAFs annotated by tissue site. **(j)** Pathway enrichment analysis of marker genes in CAFs from normal colon, primary CRC, and metastatic CRC. **(k)** TFs differentially enriched in marker peaks associated with CAFs from primary CRC, and different metastatic lesions.

By testing the relative abundance of individual stromal identities between tumours characterized by Hypercanonical and Hypocanonical transcription factor binding programs, we detected a moderate, but statistically significant higher prevalence of crypt fibroblasts, and *vice versa* a greater prevalence of pericytes in the latter (**Fig. 5e; Extended Data Fig.10a**). As crypt fibroblasts have well known trophic functions supporting intestinal self-renewal, and pericytes have a key role in tissue regeneration and angiogenesis, such findings are consistent and complementary with the corresponding tumour-intrinsic transcription factor binding programs. Additionally, tumours with an Hypocanonical transcriptional programs display a trend towards B cells depletion, as well as a higher proportion of stress-activated T cells (**Extended Data Fig.10b, c**), suggesting a convergent response to scarce microenvironmental resources.

To investigate the coordinated rewiring of the transcriptome and epigenome in TME cell populations during colorectal cancer progression, we first assessed changes in the immune and stromal composition across the normal tissue–primary tumour–metastasis continuum (**Extended Data Fig.10d-g)**. As the most abundant immune population across normal colon, primary CRC, and metastatic samples, we further characterized macrophages across these three stages (**Fig 5f**). Differential expression and functional enrichment analysis (|log2 fold change| > 1, adjusted p-value < 0.01) revealed a progressive transcriptional reprogramming along disease progression (**Fig. 5g**). In normal colon, macrophages displayed a homeostatic and antigen-presenting profile. In primary tumours, macrophages were enriched for pathways related to cytoskeletal dynamics, ECM organization, and *IL-10* signalling, indicating a shift toward a more migratory and immunomodulatory state. This phenotype was further amplified in metastatic lesions, where macrophages exhibited strong enrichment for metabolic and pro-tumorigenic pathways, consistent with adaptation to hypoxic and nutrient-deprived microenvironments **(Fig. 5g)**. To investigate the regulatory mechanisms underlying macrophage polarization, we next analysed differentially accessible chromatin regions in macrophages from normal colon, primary CRC, and metastatic samples, followed by transcription factor motif enrichment analysis (**Fig. 5h**). In all three sites, we observed strong enrichment of core macrophage lineage transcription factors, including *SPI1* (*PU.1*), *SPIC*, *MAF*, *CEBPA*, and *CEBPB*, confirming preservation of macrophage identity across disease stages. In primary CRC, macrophages additionally exhibited enrichment of *KMT2D*, *BACH1*, *BACH2*, and *CENPBD1*, indicating extensive chromatin remodelling and enhancer activation associated with immune and stress-responsive regulatory programs. In metastatic samples, this regulatory shift was further amplified by specific enrichment of *NFE2*, *NFE2L1*, and *AP-1* family members (*FOS*, *FOSL2*, *JUNB*), consistent with activation of oxidative stress, hypoxia, and inflammatory signalling pathways. Together, these results suggest a progressive epigenetic reprogramming of macrophages from a stable lineage state in normal tissue toward activated and stress-adapted regulatory states in primary and metastatic CRC, in line with the transcriptomic polarization observed along disease progression.

While several stromal populations were detected across all tissue types, CAFs were almost exclusively observed in tumour samples and were markedly expanded in metastatic lesions compared to primary tumours (**Expanded Data Fig.10f-g**). Given their tumour-specific enrichment, we focused subsequent analyses on CAFs. Examination of CAF chromatin accessibility profiles revealed clear segregation according to lesion of origin, with primary tumour CAFs clustering separately from those derived from liver (two liver lesions were assigned as Liver 1 and Liver 2) and peritoneal metastases (**Fig. 5i**). To investigate the biological basis of CAF heterogeneity across tumour sites, we performed pathway enrichment analysis on differentially expressed genes in CAFs from the primary tumour, two liver metastases, and a peritoneal metastasis (**Fig. 5j**). While all CAF populations retained core fibroblast features, including extracellular matrix organization, each lesion displayed a distinct regulatory landscape. Primary tumour CAFs were enriched for pathways associated with RUNX-dependent transcriptional regulation. In contrast, metastatic CAFs exhibited site-specific enrichments, including smooth muscle contraction and laminin interactions in Liver 1, stress-response and secretory phenotypes in Liver 2, and inflammatory signalling pathways in the peritoneal metastasis. Notably, several immune- and stress-associated pathways, including interleukin signalling, JAK–STAT signalling, and unfolded protein response programs, were preferentially enriched in metastatic CAFs, highlighting substantial functional diversification of CAF states across metastatic sites.

To identify transcriptional regulators associated with site-specific CAF chromatin states, we performed transcription factor binding site (TFBS) enrichment analysis in differentially accessible regions for each CAF population (**Fig. 5k**). Primary tumour CAFs were characterized by enrichment of developmental and mesenchymal regulators, including *TWIST1, HOX* family factors, *FOXF1, RUNX1/2/3*, and *CEBPB/D*, consistent with tissue-remodelling and fibroblast differentiation programs^31^. In contrast, metastatic CAFs displayed increased accessibility at motifs associated with immediate-early response and stress signalling pathways, including *FOSB, JUNB*, and *EGR1/3/4*, as well as chromatin architectural regulators such as *CTCF* and *E2F4*. Notably, the two liver metastases exhibited highly similar motif enrichment profiles, whereas peritoneal CAFs formed a distinct regulatory state, further supporting substantial epigenetic heterogeneity across metastatic niches. Together, these results suggest a transition from a mesenchymal, extracellular matrix–remodelling program in primary CAFs toward stress-responsive and inflammatory regulatory states extensively rewired during metastasic progression.

### CRISPRi viability screens in patient-derived-organoids (PDOs) revealed developmentally-related stress resilience dependencies

Single-cell multimodal profiling provides a granular description of cellular states and their association with clinical conditions, which can be meaningfully interpreted relying on prior knowledge, but it is inherently correlative. To begin to address causality between recurrent chromatin alterations at CRC regulatory elements and their direct impact on tumour phenotypes, we leveraged patient-derived organoids (PDOs), derived from normal, primary and metastatic CRC specimens, including 2 out 4 with matched tissue profiling by snATAC/RNAseq as described in this study (**Fig. 1b**, **Fig. 6a**). As any model only partially recapitulates real biological complexity, we compared SCAA frequency between two CRC tissue specimens (C017-T and C107-T) and their corresponding matched PDO (C017-T-PDO and C107-T-PDO). At the level of global chromatin profile, we observed between 33% (C017-T) and 43% (C107-T) of tissue SCAAs being retained in PDOs, with an overall Pearson correlation (r) between specimens and models between 0.33 and 0.46, respectively (**Fig. 6b**).

**Figure 6.**
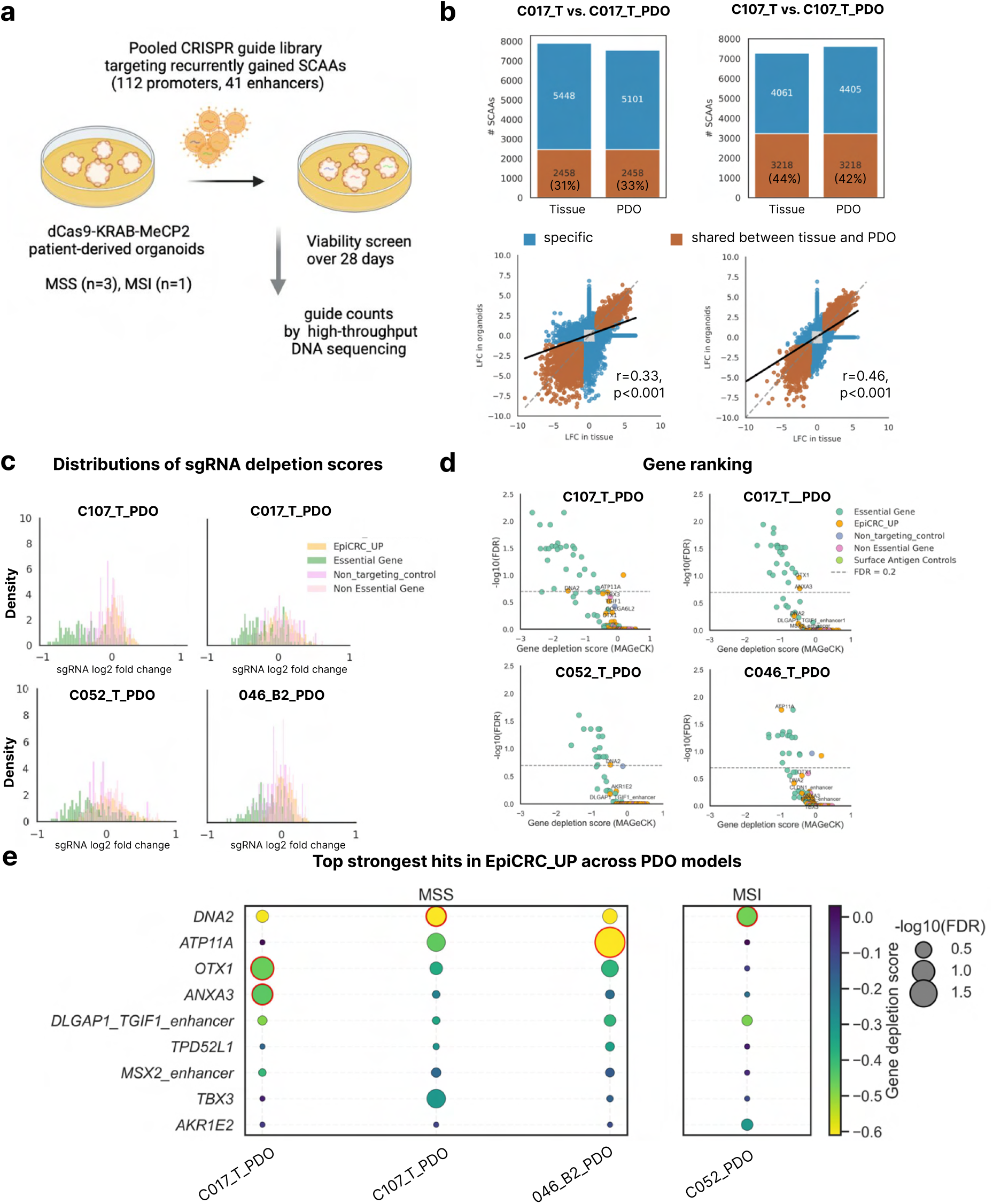
CRISPRi viability screens in patient-derived-organoids (PDOs) revealed developmentally related stress resilience dependencies. **(a)** Experimental design of pooled CRISPRi screens targeting recurrently gained SCAAs in dCas9-KRAB-MeCP2 patient-derived organoids (PDOs). **(b)** Concordance of SCAAs between C017_T_PDO and C107_T_PDO organoids and their matched tumour tissues. *Top*, number of shared and PDO-specific SCAAs. *Bottom*, Pearson correlations of log2 fold-change values between PDOs and matched tumour tissues for differentially accessible peaks. SCAAs were defined as peaks differentially accessible in tumour cells from either tissue or PDO relative to pooled normal epithelial cells (|LFC| > 1, adjusted P < 0.01).**(c)** Distribution of sgRNA depletion scores (log2 fold change) for sgRNAs targeting recurrently gained regulatory elements (EpiCRC-UP), essential genes, non-essential genes, and non-targeting controls across four PDO models. **(d)** Gene-level ranking plots from CRISPRi viability screens. Candidate dependencies associated with recurrently gained regulatory elements (EpiCRC-UP) are highlighted. Dashed lines indicate the FDR threshold of 0.2.**(e)** Summary of the strongest dependencies associated with recurrently gained regulatory elements across four PDO models. Red outlines highlight hits exceeding the significance threshold (FDR < 0.2).

To assess the impact on intrinsic tumour cell viability of the most recurrent gained SCAAs at regulatory elements, we engineered four CRC PDOs (derived from three microsatellite stable (MSS) C017-T, C046-T, and C107-T, and one microsatellite instable (MSI) C052-T CRC specimens) with a CRISPR interference (CRISPRi) chimeric epigenetic complex, based on a catalytically inactive Cas9 (dCas9) fused with the dual repressor KRAB-MeCP2^106^. We designed a custom CRISPRi guide (sgDNA) library, targeting recurrently gained regulatory elements in at least 25% of our patient cohort (occurring at 112 promoters and 41 enhancers, **Fig. 2**) with at least three independent sgRNAs. We also targeted a curated set of essential (n=52) and non-essential (n=43) genes based on well-annotated CRISPR-KO essentiality screens^107^, and included 50 non-targeting guides, for a grand total of 850 individuals sgRNAs (**Extended Data Table 7**). Upon lentiviral transduction of the sgDNA library into dCas9-KRAB-MeCP2 engineered PDOs, we recovered our cultures at regular intervals, after 2, 3 or 4 weeks and proceed to measure guide frequency across time-points by high-throughput DNA sequencing. Across models, we observed clear separation between guides targeting essential *vs.* non-essential control gene sets after three weeks in culture (**Fig. 6c**). We also found a small set of sgDNAs targeting recurrently gained SCAAs at regulatory elements (EpiCRC-UP) displaying depletion scores approaching the values observed for many known essential genes (**Fig. 6d,e**). Among those regulatory elements, we recurrently identified the promoters controlling the expression of *DNA2*, *ATP11A, OTX1*, *ANXA3*, as well as the distal enhancers associated with *TGIF1* (NC_000018.10:+:10140886-10141387) and *MSX2* (NC_000005.10:+:174520376-174520877). While DNA2, as a core component of the DNA damage response^108^ has been previously reported as common essential gene, limited dependencies were reported for the other genes in previous large-scale pan-cancer CRISPR-KO studies performed in cell lines, in agreement with previous work describing a broader dependency spectrum in PDOs^109^.

Regarding the targeted genes recurrently displaying the highest dependency scores, the flippase *ATP11A* and phospholipid-binding protein *ANXA3* are important regulators of cell membrane integrity and flexibility^110^ in response to microenvironmental stress, while *OTX1*, *TGIF1* and *MSX2* are developmentally regulated transcription factors^111–113^. Thus, we demonstrated that a small set of genes, either safeguarding cell membrane stress resilience or transcriptionally instructing developmental reprogramming, are tumour-intrinsic functional dependencies based on the viability of CRC patient-derived organoids in culture.

## DISCUSSION

Colorectal cancers are characterized by extensive genetic, transcriptional and epigenetic heterogeneity. Despite continuous progress, the precise targeting of the most common CRC genetic alterations is yet to be achieved, and the majority of patients are treated with broad spectrum chemotherapies^111^. mRNA profiling has shed light on transcriptional heterogeneity, but their interpretation is complicated by mixed cell contribution in bulk cell population assays, and the overlay between short-term and long-term sources of variations when carried out at the individual cell level as single modality^15,16,112,113^.

Here, we have performed simultaneous single-nucleus profiling of chromatin accessibility (ATAC) and 3’ mRNA expression. Across 37 CRC cases, we identify the first catalogue of transcription-affecting somatic chromatin accessibility alterations (SCAAs) that are recurrent across patients, stable between matched longitudinal samples, and target gene sets largely orthogonal to those affected by coding mutations. Promoter SCAAs converged across patients even more strongly than coding mutations or copy-number alterations, and this convergence increased from primary tumours to metastases, indicating progressive selection of a shared epigenetic landscape during CRC evolution. Approximately half of promoter SCAAs were associated with concordant changes in gene expression, rising to nearly two thirds in the presence of matched enhancer alterations. Lost SCAAs preferentially affected intestinal differentiation genes, whereas gained SCAAs targeted developmental programs, including metastasis-enriched promoter gains at *MSX2*, *PITX2*, and *TBX3*.

Transcription factor binding site analysis identified a core CRC set of inflammatory and WNT signalling-related TFs, with the former primarily driven by extrinsic oncogenic signalling, including in normal cells adjacent to tumours (*e.g*., pseudonormal), while the latter triggered by cell-autonomous genetic mechanism (*e.g*., APC^mut^). The strength of core CRC TF activity defines alternative chromatin states, termed Hypercanonical and Hypocanonical. In Hypercanonical tumours, core CRC TFs are maximally activated, in conjunction with upregulation of developmental factors binding, and repression of cell differentiation. In contrast, a translation stress-response is derepressed in Hypocanonical CRCs, dampening core CRC TF activity, and interfering with developmental reprogramming. The Hypocanonical chromatin state is particularly common but not restricted to metastases, which frequently seed microenvironments poor in nutrients, oxygen and growth factors^114,115^, as demonstrated by the depletion of trophic fibroblasts, and increased pericytes abundance, known to stimulate angiogenesis. Yet, primary CRCs are highly diverse, and a subset of them display tumour-intrinsic chromatin states, and malignant stromal cell types, reminiscent of metastases.

Moving beyond profiling chromatin alterations across CRC stages, we perturbed the most common set of promoters and enhancers recurrently gaining chromatin accessibility in more than 25% of patients in our cohort, by leveraging patient-derived organoid models and CRISPR interference. We demonstrated that a small set of genes either ensuring membrane stability and stress protection, as *ATP11A* or *ANXA3*, or programming developmental transcriptional states, like *OTX1*, *TGIF1* and *MSX2* represent novel cell viability dependencies, opening new avenues for the rational targeting of epigenetic vulnerabilities in a patient populations with urgent and unmet clinical needs.

## METHODS

### Human specimens and clinical data

Tissues and peripheral blood were obtained from patients enrolled at Fondazione IRCCS Istituto Nazionale dei Tumori, Milan. Written informed consents were obtained preceding the acquisition of the specimens. Samples and associated clinical and follow-up data were collected under an agreement for Scientific Collaboration between Fondazione IRCCS Istituto Nazionale dei Tumori and Fondazione Human Technopole (approval number 216-21).

### Nuclei extraction from clinical specimens and patient-derived organoids

Fresh frozen tissue specimens were minced into small pieces, approx. 20-50 mg in weight. A single piece was transferred into a Dounce homogenizer containing 500 μl of Nuclei extraction buffer made up of 50% Zymo Nuclei Prep buffer (cat. No. D5220) diluted as to obtain a final solution as follows: 2% nuclease-free Bovine Serum Albumin (Miltenyi Biotec, cat. No. 130-091-376), DTT 1mM, RNase inhibitors Hu 0.1 unit/μl (Qiagen, cat. No. RT-35-100), NaCl 140 mM, MgCl_2_ 3 mM, Tris-HCl pH 7.4 10 mM. Working on ice, tissues were repeatedly pestled towards homogenization for no longer than 5 minutes. At that point, tissue homogenates were diluted one-to-ten into Equilibration buffer (2% nuclease-free BSA, DTT 1mM, RNase inhibitors Hu 0.1 unit/μl, NaCl 140 mM, MgCl_2_ 3 mM, Tris-HCl pH 7.4 10 mM) and flow-through a 30 μM cell strainers (Miltenyi Biotec, cat. No. 130-110-915) to exclude aggregates and debris, followed by centrifugation at 700 g for 7 minutes with a swing bucket centrifuge refrigerated to 4℃. After supernatant removal, nuclei pellets were resuspended into 950 μl of Equilibration buffer supplemented with 50 μl of anti-nucleus microbeads (Miltenyi Biotec, cat. No. 130-132-997) and incubated for 15 minutes on ice. Upon column equilibration, nuclei-microbeads suspensions were further diluted into Equilibration buffer to reach a 2.5 ml volume, and transferred into LS columns (Miltenyi Biotec, cat. No. 130-042-401), whose ferromagnetic particle matrix retains microbead-labeled nuclei when fitted into a magnetic separator, like the Quadromacs (Miltenyi Biotec, cat. No. 130-090-976), as the solutions naturally flow through due to gravity. After washing the column with 2 ml of Equilibration buffer, the magnetic separator was disengaged, and nuclei eluted in 1 ml of Diluted Nuclei buffer (10x Genomics concentrated nuclei buffer, cat. No. PN-2000153, at 1x, DTT 1mM, RNase inhibitors Hu 1 unit/μl). Eluted nuclei were concentrated by centrifugation at 700 g for 7 minutes with a swing bucket centrifuge at 4℃, into approximately 50-100 μl. Upon staining a 5 μl aliquot one-to-one with 0.4% Trypan Blue solution (Thermo Fisher Sci., cat. No. 15250061), nuclei were visually inspected using a standard haematocytometer chamber using a bright-field microscope, followed by automated nuclei counting. Based on counts, final volume was adjusted to reach the target concentration of 4000 nuclei/μl before proceeding with the 10x Genomics Chromium Next GEM Single Cell Multiome ATAC + Gene Expression protocol, as described below. Patient-derived organoids were processed using a similar protocol, except for the initial steps, as the homogenate solutions were not obtained using a Dounce homogenizer, but by gentle pipetting of the organoid pellet.

### 10x multiome: library preparation and sequencing

Single-cell libraries were generated by following the 10x Genomics Chromium Next GEM Single Cell Multiome ATAC + Gene Expression protocol (CG000338). Briefly, nuclei suspensions were incubated with the specific transposase, able to fragment DNA in open chromatin regions and to add the adapter sequences to the ends of the DNA fragments. The transposed nuclei were loaded onto Chromium Next GEM Chip J (10x Genomics) with partitioning oil and barcoded single-cell gel beads. ATAC libraries and gene expression libraries were then prepared separately. The quantity and quality of full-length 10x cDNA, ATAC and 3′ gene expression libraries were assesed by Qubit dsDNA HS assay (Invitrogen, Q32854) and TapeStation runs (Agilent). Libraries were sequenced on Illumina platforms following 10X Genomics instructions.

### Single cell multiome samples: data processing and filtration

The sequencing short-read data generated from the Illumina platform were processed using Cell Ranger ARC^116^ (v2.0.2) for alignment to the hg38 reference genome, barcode counting, peak calling, and generation of feature-barcode matrices for both GEX and ATAC modalities. Downstream analyses, including cell and feature filtering and cell annotation, were performed on a per-sample basis using the Python packages Scanpy^117^ (v1.10.2), Muon (v0.1.6), and SnapATAC2 (v2.6.0).

For the GEX modality of each single-cell dataset, cell barcodes were filtered to retain high-quality cells (n_uniq_genes > 300, mitochondrial gene fraction < 10%), and putative doublets were identified and removed using Scrublet^118^. Gene expression counts were normalized per cell by total counts and log-transformed. The top 3,000 highly variable genes were selected using the Seurat-style dispersion-based method. Principal component analysis (PCA) was performed on scaled highly variable genes using svd_solver = ‘arpack’. Cell–cell neighborhood graphs were constructed using the first 30 principal components with 15 nearest neighbors, followed by Leiden clustering across resolutions ranging in samples from 1 to 3. Top gene expression markers in each cluster were identified using *rank_genes_groups* Scanpy function with Wicoxon rank-sum.

Initial cell type annotation was performed on a per-sample basis using the scRNA-seq modality with CellTypist framework (v1.6.3) and appropriate pre-trained models. Specifically, *Cells_Intestinal_Tract* models were used for distant normal colon samples; *Human_Colorectal_Cancer*, *Immune_All_High, Immune_All_Low*, and *Cells_Intestinal_Tract* for primary tumour samples; *Immune_All_High* and *Immune_All_Low* for lymph node samples; *Human_Lung_Atlas* for lung metastases; and *Healthy_Human_Liver* for liver metastases. Automated majority-voting annotations were subsequently refined by manual curation, assigning a consensus cell type to each Leiden cluster based on the expression of canonical marker genes.

The ATAC modality was analyzed on a per-sample basis in Python using the Muon^119^ (v0.1.6), and SnapATAC2^120^ (v2.6.0) packages. High-quality cells were retained by filtering for >2,000 unique ATAC fragments and nucleosome signal < 2, and putative doublets were removed using Scrublet. For each sample, a tile matrix was constructed using 500-bp genomic bins containing ATAC counts. A total of 25,000 most accessible features were selected with *snapatac2.pp.select_features()* with default parameters. Spectral dimensionality reduction using Laplacian Eigenmaps was performed using 30 components with *snapatac2.tl.spectral*. UMAP embeddings were computed with default settings, and a k-nearest neighbor graph was constructed using 15 neighbors, followed by Leiden clustering at a resolution of 2. Cell type annotations derived from the GEX modality were subsequently transferred to the ATAC dataset.

### Annotation of tumour cells in single-cell datasets by copy-number variation inference

To identify malignant tumour cells, we inferred copy-number variations (CNVs) as a proxy for large-scale genomic alterations. Two complementary strategies were employed for CNV inference in single-cell datasets: 1) Reference-based CNV inference using inferCNVpy (v0.5.0; https://github.com/icbi-lab/infercnvpy), which compares transcriptomic profiles of putative tumour cells against pooled non-malignant reference cells; and 2) Unsupervised CNV inference integrating both GEX and ATAC modalities using CONGAS^121^ (v1.2), which uses genome-wide segmentation input and estimates segment-level ploidy at single-cell resolution.

For the reference-based approach, inferCNVpy was applied to the GEX modality of each sample using a window size of 500 genes. Non-malignant reference cells were defined as stromal, immune, and myeloid populations based on CellTypist annotations, supported by the expression of canonical marker genes. For each cell, an L1-averaged CNV score was computed across all genomic regions. Putative tumour populations were initially identified as Leiden clusters previously annotated as epithelial cells by CellTypist, or displaying mixed cell-type annotations, and exhibiting the highest average CNV scores.

To validate tumour clusters using both GEX and ATAC modalities, we applied the CONGAS framework, which employs a Bayesian model to project single-cell RNA and ATAC profiles onto a latent space of copy-number clones. Compatible tibble objects were generated for high-quality cells from both ATAC (peak count matrices derived from Cell Ranger ARC output) and RNA (raw gene count matrices) data using *create_congas_tibbl*e(). A sample-specific genomic segmentation file was used as an input for CONGAS model obtained from ichorCNA analysis of matched low-pass whole-genome sequencing (lpWGS) data (see below). The CONGAS multiome model was initialized with negative binomial likelihoods for resulting tibble object, after excluding chromosomes X and Y. Genomic segments with weak coverage were excluded by applying non-zero cell thresholds (≥300 non-zero cells per segment per modality). Model hyperparameters were optimized using *auto_config_run()* across K = 1–10 clusters, employing diploid priors and a shrinkage parameter (λ = 0.5) to balance RNA and ATAC contributions. Final model inference was performed using *fit_congas()* (5,000 iterations, *learning rate* = 0.01, *temperature* = 20), with Bayesian Information Criterion (BIC)-based model selection and *normal*_cells = TRUE. This procedure yielded posterior estimates of segment-level copy-number states for each cluster and maximum a posteriori (MAP) cell-to-cluster assignments. ATAC Leiden clusters overlapping with GEX clusters exhibiting the highest CNV scores and corresponding to CONGAS-inferred CNV profiles consistent with WGS-derived genomic segmentation, were annotated as tumour clusters.

### Single-cell multiome cohort data integration

Raw GEX count matrices were aggregated across 33 primary tumour samples, 11 metastatic samples, and 7 distant normal colon samples. Only cells that passed quality-control filtering in per-sample analyses were retained. Mitochondrial genes and *MALAT1* were excluded from downstream analyses. Gene expression counts were normalized to a total of 10,000 reads per cell and log-transformed. 3,000 highly variable genes were re-identified using sample identity as a batch key. Principal component analysis (PCA) was performed on highly variable genes using svd_solver = ‘arpack’. To mitigate technical batch effects, Harmonypy (v. 0.0.10) was applied to the PCA embeddings using sample identity as the integration variable. Cell–cell neighborhood graphs were constructed using the first 50 Harmony-corrected PCs with 15 nearest neighbors, followed by UMAP embedding (min_dist = 0.05) and Leiden clustering at a resolution of 1.

For the ATAC modality, 500-bp tile matrices from high-quality cells were concatenated across samples. A total of 25,000 highly variable features were selected, and spectral embedding with 30 components was computed using *snapatac2.tl.spectral*. UMAP coordinates were calculated with default parameters, and a k-nearest neighbor graph with 15 neighbors was used to perform Leiden clustering at a resolution of 2. Cell type annotations derived from per-sample analyses were further refined using ATAC Leiden clustering for all cell types except tumour cells.

### Transcriptional state scoring and clustering of epithelial cells

To characterize epithelial cell states in tumour cells and pseudonormal epithelium, we curated 38 published colorectal cancer and intestinal epithelial gene signatures representing regenerative, stem-like, proliferative, differentiation, stress-response, and lineage-specific programs from previous studies^8,13,22,36,46,122–128^. Gene set activity was quantified for each cell using Scanpy’s *score_genes()* function, considering only genes detected in the dataset and using normalized expression values (‘use_raw=False”). This yielded a cell-by-signature matrix comprising 38 signature scores per cell. Signature scores were z-score standardized across cells using the *StandardScaler*() implementation in scikit-learn (v1.3). Unsupervised clustering was performed on the standardized signature score matrix using K-means clustering. The optimal number of clusters was determined by evaluating cluster solutions with 2–14 clusters using both the elbow method (within-cluster sum of squares) and the Calinski–Harabasz criterion. Based on these metrics, a three-cluster solution was selected. Final cluster assignments were obtained using K-means clustering (‘n_clusters = 3’, ‘n_init = 10’, ‘random_state = 0’). Clusters were subsequently annotated according to their dominant transcriptional programs as Stress-reactive, Oncoproliferative (ISC-like), and Oncoregenerative states.

### Differential expression analysis

For differential expression analysis (DEA), pseudobulk expression profiles were generated using the *pseudobulk*() function from decoupleR^129^ (v2.1.1) by aggregating raw gene expression counts from tumour cells or pseudonormal cells of each sample and from pooled epithelial cells derived from distant colon tissue. Cells were grouped to ensure a minimum of 80 cells per pseudobulk sample, with an approximately equal number of pseudobulk replicates generated for each tumour sample and its corresponding pooled epithelial control. Sample C072_T was excluded from this analysis due to insufficient cell numbers for pseudobulk generation under this strategy.

Within each tumour versus distant normal epithelial comparison, pseudobulk samples were filtered to retain those with at least 1,000 total counts, and genes with total counts >5 across all samples were included in the analysis. DEA was performed for each tumour sample against the pseudobulked pooled epithelial controls using PyDESeq2 (v0.5.2). For genes exhibiting high Cook’s distance, indicative of potential outliers, dispersion estimates were refitted to improve robustness (*refit_cooks* = TRUE). Wald tests were used to assess differential expression between tumour and control pseudobulk samples, yielding log₂ fold-change estimates and associated p-values. Multiple testing correction was applied using the Benjamini–Hochberg procedure to control the false discovery rate (FDR). Genes with an absolute log₂ fold change > 0.6 and FDR < 0.05 were considered differentially expressed.

### Peak calling and CNV correction of ATAC signal

500bp tile matrices of ATAC counts for pooled epithelial cells from distant colon (n=8,481) were concatenated using *anndata.concat*. Peaks were called independently in this concatenated matrix for normal epithelial cells and 500 bp tile matrixes corresponding to tumour cells or pseudonormal cells of each sample, employing the *snapatac2.macs3()* function. Only fragments shorter than 120 bp were retained for peak calling to enrich for nucleosome-free regions.

To account for copy-number alterations, peak signals in tumour samples were corrected based on ploidy and copy-number estimates from Sequenza analysis of matched deep whole-genome sequencing (WGS) samples (see Methods below):

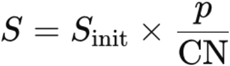

where *S*init is the initial peak signal, *p* is the estimated tumour ploidy from Sequenza, and CN represents the copy number of the segment. Inference of somatic chromatin accessibility alterations (SCAAs) with CNV correction facilitated the identification of lost SCAAs in regions with CN gain and gained SCAAs in regions with CN loss (**Supplementary Fig. 1**).

Peaks located on chromosomes X and Y were excluded. For each sample, the 20,000 top peaks by normalized signal were retained for tumour cells and pooled epithelial cells from distant colon. Peaks from all tumour samples and normal epithelial cells were then merged using an iterative process (*snapatac2.tl.merge_peaks*) to resolve overlapping peaks, yielding a final list of 91,844 non-overlapping peaks. Among these, 67,199 peaks present in at least two samples were selected for downstream analyses.

### Differential accessibility test

Prior to differential accessibility testing, ATAC-seq reads corresponding to final peaks in 500-bp tile matrices from tumour cells were corrected in each sample according to:

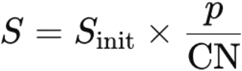

where Sinit represents the initial peak reads, *p* is the tumour ploidy estimated with Sequenza, and CN is the copy number of the segment.

Regression-based differential accessibility testing was performed using *snapatac2.tl.diff_test*, comparing each tumour sample against pooled normal epithelial cells. CN-corrected peak signals with an absolute log2fold change in RPM-normalized reads greater than 0.25 and a minimum 5% recurrence of the peak in both groups were retained for analysis. To assess whether cell-type labels provide predictive information about peak presence beyond coverage-based features, we performed a likelihood ratio test using logistic regression models. For each genomic region, a reduced model was fit using only per-cell coverage features, and a full model included both coverage features and one-hot encoded cell-type labels. Both models were fit without regularization using the *lbfgs* solver, and log-likelihoods were computed from the negative log-loss. The test statistic was defined as:

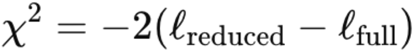

where *ℓ*_full_ is log-likelihood of full model (coverage + cell-type label), and *ℓ*_reduced_ is log-likelihood of reduced model (coverage only). Under the null hypothesis that the additional features do not contribute to prediction accuracy, the likelihood-ratio statistic is asymptotically chi-squared distributed, with degrees of freedom corresponding to the difference in the number of parameters between the full and reduced models (*df* = *p*_1_ − *p*_O_). P values were derived from the chi-squared survival function. Significant tests were interpreted as evidence that cell-type labels provide predictive information beyond coverage alone. Multiple hypothesis testing was controlled using the Benjamini–Hochberg false discovery rate procedure.

Differentially accessible peaks in tumour cells or pseudonormal cells relative to normal distant epithelial cells (|log_2_ fold change| > 1, adjusted p < 0.01) were retained for annotation of somatic chromatin accessibility alterations (SCAAs).

### Annotation of peaks and definition of SCAAs

Gene annotations from the GENCODE evidence-based annotation of the human genome (GRCh38), release 32 (Ensembl 98) were used to assign genomic features corresponding to differentially accessed peaks in every tumour or pseudoepithelial sample. Promoter regions were defined in a strand-aware manner: for transcripts on the “+” strand, promoters encompassed the interval from 1000 bp upstream to 500 bp downstream of the transcription start site (TSS), whereas for transcripts on the “−” strand, promoters were defined as the region extending from 500 bp upstream to 1000 bp downstream of the TSS. To annotate enhancers genomic regions corresponding to enhancers from GeneHancer_v5.22 database were chosen. In **Fig. 1f** and **Extended Data Fig. 6a** chromatin regions were annotated as enhancers based on overlap with candidate cis-regulatory elements (cCREs) from the ENCODE Registry of Candidate Cis-Regulatory Elements derived from male and female transverse colon tissues. Peaks overlapping ENCODE enhancer-like signatures classified as proximal enhancer-like (pELS), distal enhancer-like (dELS), pELS,CTCF-bound, or dELS,CTCF-bound were designated as enhancers.

Differentially accessible peaks overlapping promoters and enhancers were annotated as somatic chromatin accessibility alterations (SCAAs) only if the expression of the corresponding gene was also differentially regulated between tumour and distant normal cells (see Differential Expression Analysis, Methods). For matched samples from the same patient (n pairs = 7), SCAAs were retained only if detected in both samples.

### Similarity of somatic chromatin accessibility alteration (SCAA) in matched and unmatched CRC samples

The cohort included seven pairs of matched samples from the same patient, comprising paired primary tumours and distant metastases (n = 1 pair), multiple distant metastases from distinct organs (n = 1 pair), and paired pre- and post-treatment primary tumours (n = 5 pairs). To assess the similarity of SCAA profiles between colorectal cancer (CRC) samples, differentially accessible peaks between tumour and normal cells (|log2 fold change| > 1, adjusted p < 0.01) were selected, and Spearman’s correlation coefficients were calculated between pairs of samples using log_2_ fold change values. Genomic regions with |log2fold change| < 1 or adjusted p > 0.01 were assigned a value of 0.

To estimate the proportion of shared SCAAs across matched and unmatched samples, the number of shared SCAAs defined as regions with |log2 fold change| > 1 and concordant direction was calculated pairwise and compared between matched and unmatched sample groups.

### Transcription Factor Motif Enrichment and Clustering of TFs

Motifs corresponding to transcription factor (TF) binding sites were assessed in differentially accessible peaks using a dataset of binding sites for 1,165 TFs from CIS-BP database^130^. Motif enrichment was calculated with *snapatac2.tl.motif_enrichment* in upregulated tumour peaks or pseudonormal epithelial cells (log_2_ fold change > 1, adjusted p < 0.01 for tumour cells; log_2_ fold change > 1, adjusted p < 0.01 for tumour-adjacent non-malignant epithelial cells) relative to peaks from distant normal epithelial cells, which were used as background (log_2_ fold change < -1, adjusted p < 0.01.

A total of 821 significantly enriched TFs (|log_2_ fold change| > 1, adjusted p < 0.01) that were recurrent in at least five samples were selected. These TF enrichment values were standardized, and dimensionality was reduced via principal component analysis (PCA), retaining two principal components to represent both normal and tumour samples in the space of TF activity.

A total of 361 significantly enriched TFs (|log_2_ fold change| > 1, adjusted p < 0.01) that were recurrent in at least seven tumour samples were selected for downstream clustering to identify TF groups based on their representativeness across tumour samples. To identify robust and stable clusters of transcription factors (TFs), we implemented a bootstrap consensus clustering approach on a matrix of enrichment signals for recurrent TFs. Briefly, for each number of clusters *k* from 3 to 25, 80 bootstrap resamples were generated by randomly selecting 80% of TFs without replacement. Each resampled matrix was clustered using hierarchical clustering with Ward linkage and Euclidean distance, and a consensus matrix was updated to reflect the frequency with which pairs of TFs co-clustered across bootstraps. The consensus matrix was then normalized by the number of bootstrap iterations, and cluster stability was computed as the average pairwise consensus within clusters derived from the full dataset. Stability scores were used to evaluate the robustness of clustering across different values of *k*, and *k*=4 was selected for the final clustering. Clusters were annotated according to TF families, and cluster-specific signals were compared between primary tumours and metastases.

Scaled values of cluster-specific TF enrichment per sample were used as input for dimensionality reduction using UMAP with 5 nearest neighbors, a minimum distance of 0.3, and Euclidean distance as the metric. The resulting UMAP coordinates were used to visualize similarities between samples based on their TF cluster enrichment profiles.

Samples were further stratified into two groups based on their TF enrichment profiles by cutting the hierarchical clustering dendrogram at k = 2 using the SciPy fcluster function (criterion = “maxclust”). To identify transcriptional programs recurrently activated within TF-defined patient groups, differential expression results (see ‘Differential expression analysis’ above) were binarized using thresholds of |log2 fold change| > 1 and adjusted P < 0.01. For each gene, recurrence was calculated as the proportion of samples within a group in which the gene was significantly upregulated. Genes recurrently upregulated in at least 50% of samples in the group were identified. Group-specific recurrent genes were defined as those showing a recurrence frequency difference of at least 20% between groups while remaining recurrent (≥50%) in the group of interest.

### Single-cell transcription factor activity inference

TF activity was inferred from single-cell ATAC-seq data for a subset of patients (002_S1, 006_S1, 008_S1) where all three types of epithelial cells existed (normal and pseudonormal epithelium, tumour cells) using chromVAR implemented through the pychromVAR package^131^ (v.0.0.4). ATAC fragments from the selected subset of multiome samples were imported into SnapATAC2 and processed to generate a unified peak-by-cell accessibility matrix. Peaks were called separately for each annotated cell type using MACS3 and merged into a consensus peak set. DNA sequences underlying accessible regions were retrieved from the hg38 reference genome, and GC-content bias was computed to generate matched background peaks. TF motifs from the JASPAR 2020 CORE vertebrate collection were mapped to accessible regions using pychromVAR. chromVAR deviation scores, representing motif accessibility relative to GC-matched background expectations, were then calculated for each cell and motif. The resulting deviation matrix was integrated with the corresponding RNA annotations and low-dimensional embeddings, enabling visualization and comparison of TF activity across epithelial cell states.

### Analysis of stromal and immune cell subsets

To characterize stromal and immune cell heterogeneity, cells initially annotated as stromal or immune were extracted from the integrated single-cell dataset and reanalyzed independently. In RNA modality raw UMI counts were normalized to 10,000 counts per cell, log-transformed, and highly variable genes were identified using Scanpy. Principal component analysis (PCA) was performed on the top 3,000 highly variable genes, followed by batch correction across patients using Harmony. A nearest-neighbor graph was constructed using the Harmony-corrected principal components, and cells were clustered using the Leiden algorithm. Uniform Manifold Approximation and Projection (UMAP) was used for visualization of stromal cell states. Cell identities were assigned based on cluster-specific marker genes identified using the Wilcoxon rank-sum test implemented in *scanpy.rank_genes_groups()* together with canonical lineage markers. In stromal cells this approach resolved endothelial cells, lymphatic endothelial cells, pericytes, myofibroblasts, cancer-associated fibroblasts (CAFs), WNT5B-positive fibroblasts, adventitial fibroblasts, progenitor fibroblasts, glial cells, and a cluster of contaminating cells that were subsequently removed from the analysis. In immune subset B and plasma cells, CD4+, CD8+, cycling, migratory, stress-activated and regulatory T cells, dendritic cells, mast cells, and macrophages were identified.

To validate transcription factor programs identified within stromal and immune subpopulations, pseudobulk chromatin accessibility profiles were generated separately for each annotated cell type by aggregating single-cell ATAC-seq fragments across cells from different patients. Peak calling was performed on each pseudobulk profile using MACS3 (see details above), and marker peaks were identified between stromal populations using *snapatac2.tl.marker_regions*() (P<0.05). Enrichment of transcription factor motifs within cell type-specific accessible regions was assessed using *snapatac2.tl.motif_enrichment()* and validated by chromVAR deviation scores at single-cell resolution, enabling the identification of transcription factors associated with distinct stromal states.

We next investigated chromatin accessibility differences in CAF and macrophage populations derived from primary tumours and metastatic lesions. CAFs and macrophages were stratified according to tissue origin and aggregated into group-specific pseudobulk ATAC-seq profiles. Peaks were called independently for primary tumour-derived and metastasis-derived populations using MACS3, and differential accessibility analyses were performed to identify regions exhibiting group-specific chromatin accessibility. Transcription factor motif enrichment analyses were subsequently conducted on differentially accessible regions in genome-wide background model to identify candidate regulatory factors associated with metastatic CAF and macrophage states.

### Bulk Whole-Genome Sequencing: Extraction, Library Preparation and Data Generation

gDNA from blood, tissues and PDOs for bulk characterization were extracted by using the QIAamp DNA Blood Mini Kit (Qiagen, catalog no. 51104), the QIAamp DNA Mini Kit (Qiagen, catalog. no. 56304) and the AllPrep DNA/RNA Mini Kit (Qiagen, catalog no. 80204), respectively. gDNAs were used as input to generate whole genome libraries by using NEBNext Ultra II FS DNA Library Prep kit for Illumina (New England Biolabs, catalog no. E6177). The quality of the libraries was checked by D1000 ScreenTape (Agilent, catalog no. 5067-5582) on a 4200 TapeStation System (Agilent). Whole genome sequencing (WGS) on pooled whole genome libraries was performed on Illumina platforms (150 paired-end reads).

### Bulk Whole-Genome Sequencing: Data Processing

The FASTQ files generated from deep WGS were aligned to the human reference genome (GRCh38) using the Burrows-Wheeler Aligner (BWA) algorithm^132^. Duplicate reads were identified and marked using Picard’s Mark Duplicates tool. Somatic mutations were then called using Mutect2 (v4.1.0.0), with the tumour sample compared against a peripheral blood monocyte DNA control. Sequenza (v2.2.0; https://bitbucket.org/sequenzatools/sequenza/src/chemins/) was used for cellularity and ploidy estimation and copy number detection. For downstream analysis and tracing relationships between genome and epigenome high-quality (passed mutect2 filtration criteria) protein-affecting variants with coverage more than 10 reads in both tumour and normal samples, with at least 3 reads supporting altered allele in tumour, were selected.

### Analysis of convergence of genomic and epigenomic alterations

To compare the recurrence of genetic and epigenetic alterations across colorectal cancers, we generated gene-level alteration profiles for each tumour sample. The gene universe was defined as all genes associated with promoter peaks detected in the multiome cohort (see *Peak calling*), comprising 12,900 genes. For each tumour, three gene sets were generated: (i) genes harbouring protein-altering somatic variants (missense, nonsense, indels, translation start site or splice site mutations); (ii) genes located within copy-number gain or loss segments relative to the estimated tumour ploidy; and (iii) genes with promoter-associated somatic chromatin accessibility alterations (SCAAs) predicted to affect gene expression. Pairwise contingency tables were constructed for all tumour pairs within the primary and metastatic cohorts separately, and the degree of overlap between alteration profiles was quantified using Fisher’s exact test. Odds ratios (ORs) were calculated from the contingency tables after applying a Haldane-Anscombe correction (adding 0.5 to each cell) to avoid undefined estimates in the presence of zero counts. Because microsatellite instability (MSI) tumours exhibit a markedly distinct mutational and copy-number alteration landscape that strongly influences OR estimates, MSI samples were excluded from this analysis. Longitudinally matched samples from the same patient were also excluded to ensure independence between comparisons.

### Genotyping of somatic variants and estimation of per-cell mutational burden

Somatic variants were genotyped using cellsnp-lite (v1.2.1) on single-cell ATAC-seq BAM files, using filtered VCF files derived from matched whole-genome sequencing (WGS) data. Genotyping was performed with the following parameters: - p 2 --minMAF 0.005 --minCOUNT 1. Only high-confidence genotyped variants were retained for downstream analysis (a total coverage greater than 10 reads in both tumour and normal samples, with at least 3 reads supporting the alternative allele in tumour cells).

To mitigate the sparsity inherent to single-cell ATAC-seq data, allele counts were aggregated across all genotyped somatic variants per cell. For each cell, the total number of reads supporting the alternative allele (AD) and the total number of reads covering variant positions (DP) were summed across all variants. The per-cell mutational burden was then quantified as the AD/DP ratio, calculated as:

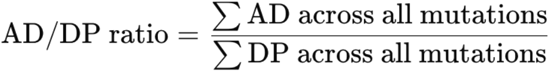

### Functional Enrichment Analysis

Functional enrichment analysis of recurrent SCAA-associated genes in tumour was performed using the STRING database (v12.0; https://string-db.org/). Gene lists were submitted using the *Multiple proteins* option for *Homo sapiens*, and enrichment of Gene Ontology (GO), Reactome, KEGG and other pathway annotations was assessed using the built-in STRING enrichment analysis. Significantly enriched terms were ranked according to the STRING statistical framework, and pathways with a false discovery rate (FDR) < 0.05 were considered statistically significant.

Functional enrichment analyses of recurrent SCAA-associated genes in pseudonormal epithelial cells, hypo- and hypercanonical TF programs, and marker genes from macrophages and CAFs were performed using the Enrichr web platform (https://maayanlab.cloud/Enrichr/). Lists of differentially expressed genes were queried against Gene Ontology (GO) Biological Process, Reactome Pathways 2024, KEGG 2021 Human, MSigDB Hallmark 2020, and Elsevier Pathway Collection databases. Enriched terms were ranked according to the Enrichr statistical framework, and pathways with a false discovery rate (FDR) < 0.05 were considered significantly enriched. For enrichment analyses of recurrent SCAAs in pseudonormal epithelial cells, an FDR threshold of < 0.1 was used because of the limited number of recurrent genes.

### Patient-derived colorectal cancer organoids (PDOs)

PDOs were established following previously published procedures. Briefly, tissue specimens were minced and digested with collagenase II (5 mg ml−1; Gibco) and dispase I (1.25 mg ml−1; Gibco) in human basal medium (Advanced Dulbecco’s modified eagles medium with nutrient mixture F-12 Hams (Gibco) supplemented with HEPES (10 mM; Gibco), Glutamax (2 mM; Gibco) and Primocin (1 mg ml−1; InvivoGen)) at 37 °C for 30-60 min, followed by an additional 5-10 min digestion with TrypLE (Gibco) at 37 °C. The digested materials were suspended in Matrigel, dispensed onto 6-well plates as 50 μL droplets, and, after Matrigel form a gel, overlaid with the following medium: human basal medium supplemented with B-27 Supplement (1X; Gibco), 10 nM gastrin I (Sigma-Aldrich), 1.25 mM N-acetylcysteine (Sigma-Aldrich), 10mM Nicotinamide (Sigma-Aldrich), 100 ng/ml recombinant mouse Noggin (PeproTech), 50 ng/ml recombinant human EGF (PeproTech), 100 ng/ml recombinant human IGF-1 (PeproTech), 10 ng/ml recombinant human FGF-basic (FGF-2) (PeproTech), 500 ng/ml recombinant human R-spondin1 (PeproTech), 500 nM A83-01 (Tocris Bioscience) and 100ng/mL of recombinant human Wnt-3a (R&D). Media were refreshed every 3–4 days. For organoid propagation, confluent organoids were removed from Matrigel, dissociated into small clusters of cells by pipetting, and resuspended in an appropriate volume of fresh Matrigel. Organoids were routinely tested for the presence of Mycoplasma contamination using the Mycoalert Mycoplasma Detection kit (Lonza).

### Lentiviral vector packaging

For lentiviral vector packaging, 3×10^5^ cells HEK293T/17 (ATCC, Catalog number CRL-11268) were plated in 2 mL DMEM supplemented with 10% FBS without antibiotics on a single-well of a standard 6-well tissue culture plate. The transfection mix was prepared as follows (reagents per well of a 6-well plate): 1.5 μg psPAX2 (Addgene #12260), 0.4 μg pMD2.G (Addgene #12259), 4 μg of transfer plasmid in 300 μL Opti-MEM (Gibco, cat. No. 31985062), 18 μg Fugene HD transfection reagent (Promega, cat. No. E2311). After 20 minutes of incubation at room temperature, the transfection mix was added slowly and dropwise to each well of HEK293T/17 cells and, after 16 hours of incubation in a standard TC-incubator, was replaced with DMEM supplemented with 10% FBS without antibiotics. After 72 hours post-transfection, lentiviral vector containing supernatant was harvested, and centrifuged at 500 g for 10 minutes at 4°C to and filtered through a 0.45 μm syringe filter into a fresh 15 mL conical tube to remove any potential cellular debris. For each lentiviral vector, viral supernatant were concentrated using Lenti-X Concentrator (Takara Bio, cat. No. 631231), according to manufacturer’s instructions. Brieflly, Lenti-X Concentrator (1 volume) were added to the viral supernatant (3 volumes), mixed by inversion and incubated at 4°C for 30 minutes. Afterwards, the suspension was centrifuged at 1,500 g for 1 hour at 4°C. Upon careful supernatant removal, a visible off-white pellet was vigorously resuspended in 300 ul OptiMEM, and stored at –80°C for subsequent usage.

### Generation of dCas9-KRAB-MeCP2 expressing patient-derived organoids

To generate dCas9-KRAB-MeCP2 expressing patient-derived organoids (PDOs), PDOs were gently dissociated to single cells using standard cell culture procedures and counted. Approximately one million single cells were resuspended in CRC media, supplemented with ROCK inhibitor (10 μM final, Sigma Aldrich, cat. No. Y0503) and polybrene (10 μg/mL; Sigma-Aldrich, cat. No. TR1003), and mixed with lentiviral particles generated using the transfer vector Lenti-SFFV-dCas9-KRAB-MeCP2-P2A-Bst (Addgene, plasmid #122205), expressing a bicistronic construct encoding for the chimeric dCas9-KRAB-MeCP2 and the blasticidin (Bst) resistance gene, under the control of the SSFV promoter. Cell suspensions mixed with lentiviral particles were transferred on non-adherent tissue-culture plates and spinoculated for 1 hour at 100 g in a swing bucket centrifuge, followed by static incubation for 1 hour in a standard tissue culture incubator. Afterwards, cells were collected by centrifugation, washed and resuspended in 500 μl of fresh Matrigel, plated as individual domes, which were left to jellifying in theincubator for 30 minutes, before being overlaid with medium. After two days, blasticidin was added at 10 μg/mL to select transduced cells over at least ten days in culture.

### CRISPR interference (CRISPRi) pooled library design and production

A custom pooled CRISPRi guide (sgRNA) library, targeting recurrently gained regulatory elements in at least 25% of our patient cohort (112 promoters and 41 enhancers, **Fig 2**) was compiled including at least three independent guides *per* targeted locus. We also targeted a curated set of essential (n=52) and non-essential (n=43) genes based on well-annotated CRISPR-KO essentiality screens^107^, and included 50 non-targeting guides, for a grand total of 850 individuals sgRNAs. Most guides targeting promoters were selected from Sanson et al. 2018^133^ (*e.g*., 680 out 850 guides). Additionally, we designed *de novo* 123 guides targeting 41 enhancers, as well as 47 guides targeting germline variants occurring at regulatory elements, as identified by WGS analysis of our patient cohort. The full guide list is provided as **Extended Data Table** 7. The lentiviral backbone vector was encoded by a proprietary plasmid pRSGScribe1-EFS-TagRFP-2A-Puro-U6-sg-3LTR (Cellecta, USA) expressing sgRNAs under the control of the U6 promoter, and a bicistronic construct including TagRFP and Puromycin resistance gene, for positive selection based on either RFP fluorescence or puromycin treatment. The library was packaged in lentiviral vectors by an external supplier (Cellecta, USA), titrated by flow cytometry, and DNA high throughput sequencing was performed to assess guide distribution by a commercial service, finding 90% of guide sequences within 1.67-fold count range, and 99% within 2.36-fold count range.

### CRISPRi screens on patient-derived organoids

Test experiments were performed to titrate the volume of lentiviral vectors, encoding the CRISPRi pooled guide library, needed to achieve an approximate multiplicity-of-infection (MOI) equal to 0.3 in patient-derived organoids (PDOs), as assessed by automated cell counting based on RFP fluorescence. Prior to set up each CRISPRi screen, PDOs were expanded in suspension cultures, including 10% v/v Matrigel in standard media, within sterile glass flask maintained under constant orbital agitation, and regularly supplemented with fresh media and resuspend by pipetting every three or four days. On the first day of the CRISPR screen, PDOs were gently dissociated to single cells, by removal of residual Matrigel using Cell Recovery Solution (Corning, cat. No. 354253), followed by gentle trypsinization with 0.5X TrypLE (Gibco, cat. No. A1217701) at 37℃ in constant agitation for an interval comprised between 30 and 60 minutes. Upon reaching extensive single-cell dissociation, as determined by visual inspection using an inverted bright-field microscope, suspensions were flowed through a 30 μM cell strainers (Miltenyi Biotec, cat. No. 130-110-915) to exclude debris, followed by centrifugation at 200 g for 5 minutes with a swing bucket centrifuge refrigerated to 4℃. Upon supernatant removal, cells were resuspended in 8 ml of CRC media, supplemented with ROCK inhibitor (10 μM final, Sigma Aldrich, cat. No. Y0503) and polybrene (10 μg/mL; Sigma-Aldrich, cat. No. TR1003), cell concentration was measured using automated cell counting, and final volume adjusted to reach the target of 3 x 10^6^ cells/ml. For each individual CRISPRi screen, 24 x 10^6^ cells/ml in 8 ml of CRC media were mixed with an adequate volume of lentiviral particles suspension, as to achieve an approximate 0.3 MOI and a coverage of 200 cells transduced for each individual guide. Each millilitre of cell suspension mixed with lentiviral particles was transferred into a single well of a non-adherent 24-MTP tissue-culture plate and spinoculated for 1 hour at 100 g in a swing bucket centrifuge with temperature set at 30℃, followed by static incubation for 1 hour in a standard tissue culture incubator. Afterwards, cells were collected by centrifugation, washed and resuspended in 48 ml of fresh CRC media, supplemented with 10% v/v Matrigel, and 10 μM of ROCK inhibitor, and plated by adding 2 ml of cell suspension into 4 x 6-MTP ultralow attachment multiculture plates (Corning, cat. No. 3471) for a total of 24 wells. After 72 or 96 hours post-transduction, puromycin was added at the final concentration of 5 μg/ml for negative selection of untransduced cells. Positively selected PDOs were collected at regular intervals after 11 or 12 days (time point 1, T1), after 21 days (T2), and after 28 days (T3) when the experiment was terminated, by harvesting 5 wells *per* time point as independent replicas. Organoids were harvested using standard procedures, and intact organoids were concentrated by centrifugation, supernatant removal, and cryopreserved at -20℃. At the end of the time course, genomic DNA extraction was performed in parallel across all cell pellets (Qiagen kit).

### Amplicon DNA sequencing of pooled CRISPRi guide libraries

Genomic DNA (gDNA) was quantified using both a Nanodrop spectrophotometer and a Qubit analyser using the dsDNA BR Assay Kit (Invitrogen, cat. No. Q32850). For PCR amplification, a first PCR was performed using 3.3 μg of gDNA for each replica mixed with 1x NEB Next Ultra II Q5 Master Mix (New England Biolabs, cat. No. M0544L), and 1 μl of combined forward and reverse primers at 10 μM each, in a final volume of 100 μl of total volume. Cycling conditions were as follows: an initial 30 sec at 95 °C; followed by 10 sec at 95 °C, 75 sec at 65 °C, for 17 cycles; and a final 10 min extension at 65 °C. Out of this total volume, 25 ul were utilized as template for a second PCR using standard Illumina indexing primers. PCR products were purified with Agencourt AMPure XP SPRI beads according to manufacturer’s instructions (Beckman Coulter, A63880). Libraries were sequenced on DNA high throughput sequencer NextSeq 2000 (Illumina), loaded with a 5% spike-in of PhiX DNA, using either a single-end or pair-end protocol with at least 100 cycles, with a minimal target of 5 million single reads/sample. Primers are reported in **Extended Data Table 7**.

### CRISPRi screen analysis

Sequencing reads from pooled CRISPRi screens were processed using MAGeCK^134^ (v. 0.5.9.5) to quantify sgRNA abundances. Samples with at least 7M total sequencing depth, 85% mapped reads, 0 undetected sgRNAs, Gini index <0.1 were considered as passed QC. sgRNA count matrices were generated for all samples and merged into a single dataset for downstream analysis.

To compare guide abundance across samples, sgRNA counts were normalized by the median count of each sample and transformed into log2 fold changes relative to the plasmid library reference. Screen performance was evaluated using predefined control guide categories, including essential genes, non-essential genes, and non-targeting controls. Distributions of sgRNA fold changes were examined across biological replicates and experimental conditions, and receiver operating characteristic (ROC) analyses were performed to assess the ability of guide depletion to distinguish positive and negative control sets. Samples with ROC AUC for essential set of genes>0.8 were considered as passed depletion score QC.

Gene-level enrichment and depletion statistics per sample were calculated with MAGeCK run command relative to the plasmid library reference. Genes targeted by multiple sgRNAs showing consistent negative depletion relative to the plasmid library and passing the selected FDR<0.2 threshold were considered significant hits.

### Statistical analysis and visualization

Comparisons between two independent groups of samples (e.g., primary versus metastatic tumours) were performed using the Mann–Whitney U test. P-values from multiple pairwise comparisons were adjusted using the Benjamini–Hochberg false discovery rate (FDR) procedure. Statistical significance was defined as adjusted *P* < 0.05. Comparisons between paired samples (e.g., numbers of gained and lost SCAAs within the same samples) were conducted using the paired Wilcoxon signed-rank test. Spearman’s rank correlation coefficient was used to assess correlations between numerical variables, such as in analyses of SCAA profile similarity across samples. All mentioned statistical tests were used as implemented in the scipy (v.1.14.1) Python package^135^. All plots were generated using matplotlib^136^ (v3.9.2) and seaborn (v0.13.2), and statistical annotations were added using the statannotations Python package (v0.7.2; https://doi.org/10.5281/zenodo.7213391). All boxplots show the median (center line), interquartile range (box), and 1.5× interquartile range (whiskers). Individual observations are shown as jittered points.

## Supplementary Material

**Extended Data Table 1.** Clinical data from 37 patients in CRC multiome cohort

**Extended Data Table 2.** a. Cell markers used to annotate main cell types b. Source papers used to extract gene signatures describing transcriptional states across epithelial cells in CRC c. 38 gene sets from 12 previous studies on CRC transcriptional heterogeneity to identify transcriptional states in epithelial cells

**Extended Data Table 3.** Top SCAAs affecting promoters and enhancers recurrent in at least 25% of patients in CRC cohort used in CRISPRi screen

**Extended Data Table 4.** a. Results of STRING-DB GSEA for recurrently gained and lost SCAAs affecting promoters in tumour samples (present in ≥9/36 patient samples). b. Results of STRING-DB GSEA for recurrently gained and lost SCAAs affecting promoters in pseudonormal cells (present in ≥2/12 patient samples). c. Results of GSEA for recurrent differentially expressed genes (DEGs) in the Hypocanonical and Hypercanonical groups of CRC samples.

**Extended Data Table 5.** a. TF motifs enriched in pseudonormal cells in CRC cohort b. TF motifs enriched in tumour cells in CRC cohort

**Extended Data Table 6.** DEGs in Hypo- and Hypercanonical groups recurrent in at least 50% or samples from the group with delta>20% compared to another group

**Extended Data Table 7.** a. Custom CRISPRi screen library structure b. 850 individual sgRNAs used in custom CRISPRi screen c. Target annotations d. Primers used in amplicon DNA sequencing of pooled CRISPRi guide libraries

## Acknowledgements

This work was supported by the Associazione Italiana per la Ricerca contro il Cancro (AIRC) to A.S. (28961). A.S. is also supported by the ERC Consolidator Award to A.S. (101125077). We thank Konstantin Winter and Anastasiia Romanova for the invaluable support to the Computational Biology Research Centre at Human Technopole. The authors would like to acknowledge Didier Trono and Andrea Califano for depositing plasmids used in this work and Addgene for plasmid distribution.

## Data availability

The raw sequencing data generated in this study have been deposited in the European Genome-phenome Archive (EGA) and will be made available through controlled access owing to patient privacy considerations. The submission is currently associated with Data Access Committee accession EGAC50000001117. Processed data, including single-cell gene expression and chromatin accessibility matrices, are archived in Zenodo (https://doi.org/10.5281/zenodo.21834761).

## Code availability

All custom code used for data processing, statistical analyses and figure generation in this study is available from the GitHub repository (https://github.com/sottorivalab/SCAA_2026).

