## Extended Data Figures for "Epigenetic evolution of colorectal cancer and its microenvironment reveals new vulnerabilities"

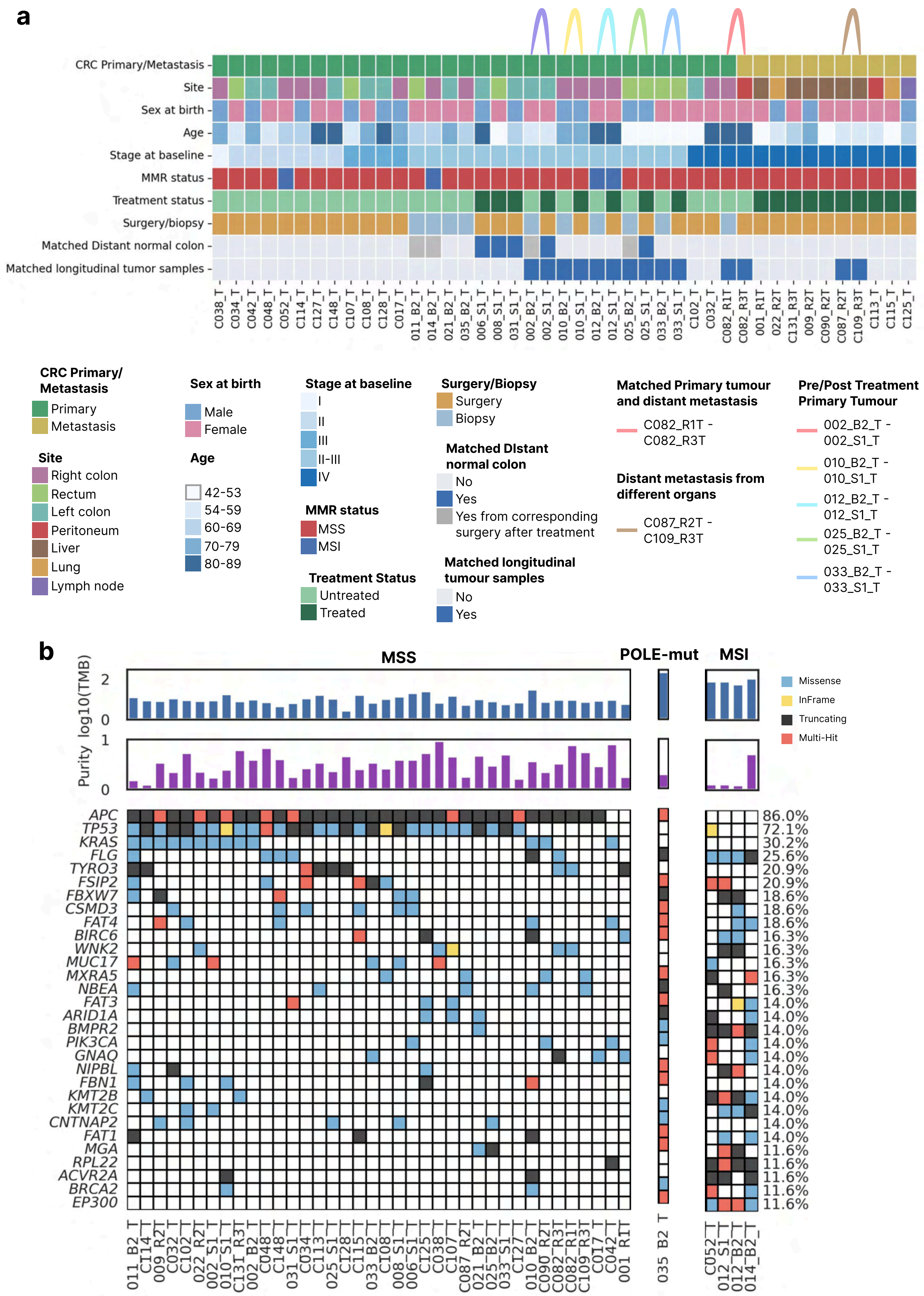


**Extended Data Figure 1.** **(a)** Clinical characteristics of biospecimens in the study cohort. **(b)** Genetic landscape of samples collected and processed with whole-genome sequencing. Tumor purity as predicted with Sequenza (see Methods) is shown.

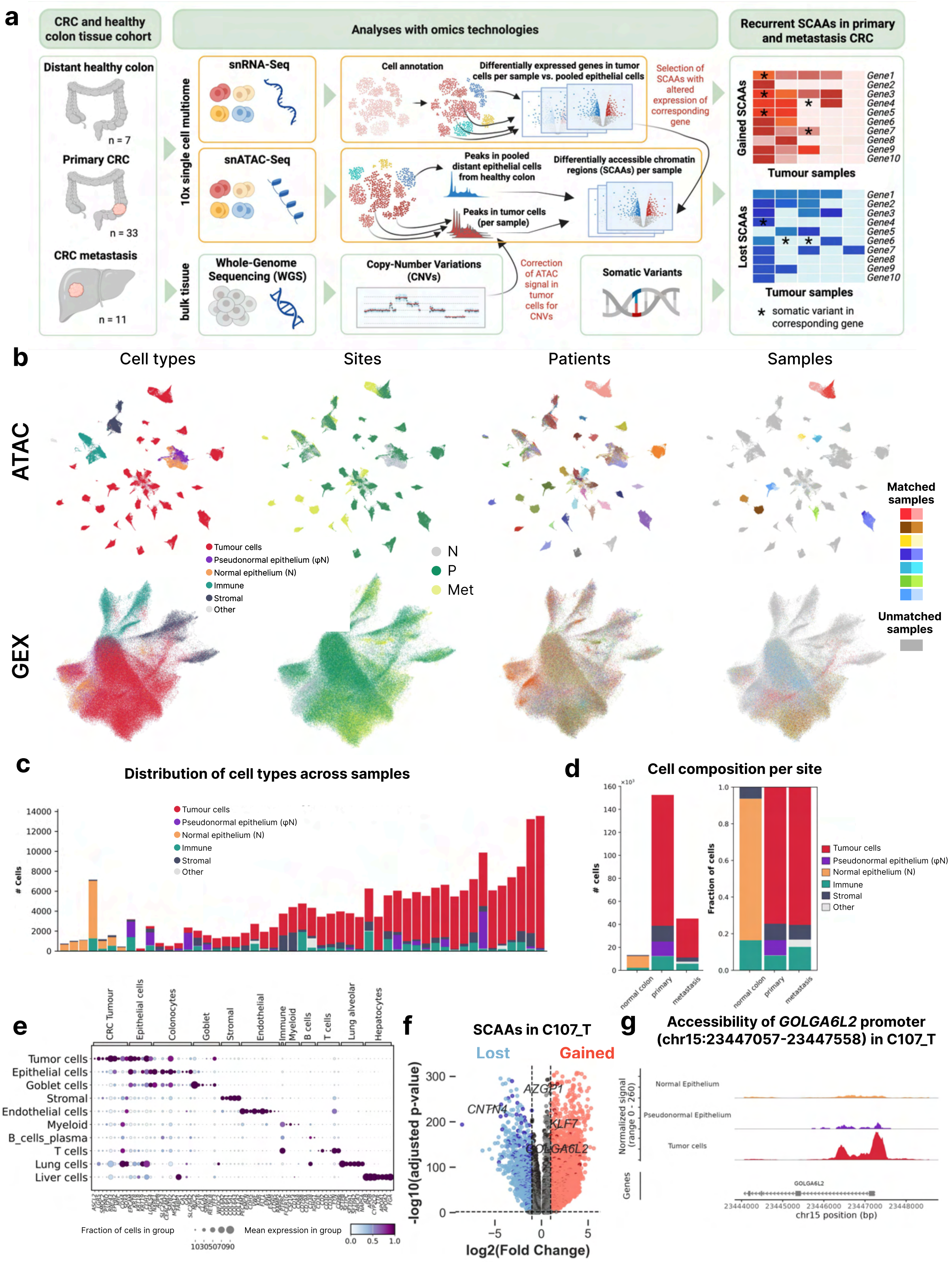


**Extended Data Figure 2. (a)** Experimental design **(b)** UMAP projections (GEX and ATAC) showing cell types, tissue site, patient ids, and matched samples from the same patient. **(c)** Sample cell composition sorted by percentage of tumor cells. **(d)** Cell composition of normal colon controls, primary CRC and metastatic samples shown as absolute cell numbers (*left*) and proportional representation (%) (*right*). **(e)** Top five marker genes expressed in each annotated cell type across the scRNA modality of multiome cohort. **(f)** Volcano plot showing differentially accessible peaks upregulated (gained SCAAs, red) and downregulated (lost SCAAs, blue) in tumor of C107_T sample against pooled normal epithelial cells. Thresholds for differentially accessible peaks are |log2foldchange|>1, padj<0.01.Dark red and bark blue dots are peaks corresponding to gene gained and lost promoters, respectively. **(g)** Track plot showing normalized ATAC-seq signal for a promoter SCAA in the *GOLGA6L2* gene in pooled normal epithelial cells from distant colon, pseudonormal epithelium, and tumor cells from sample C107_T.

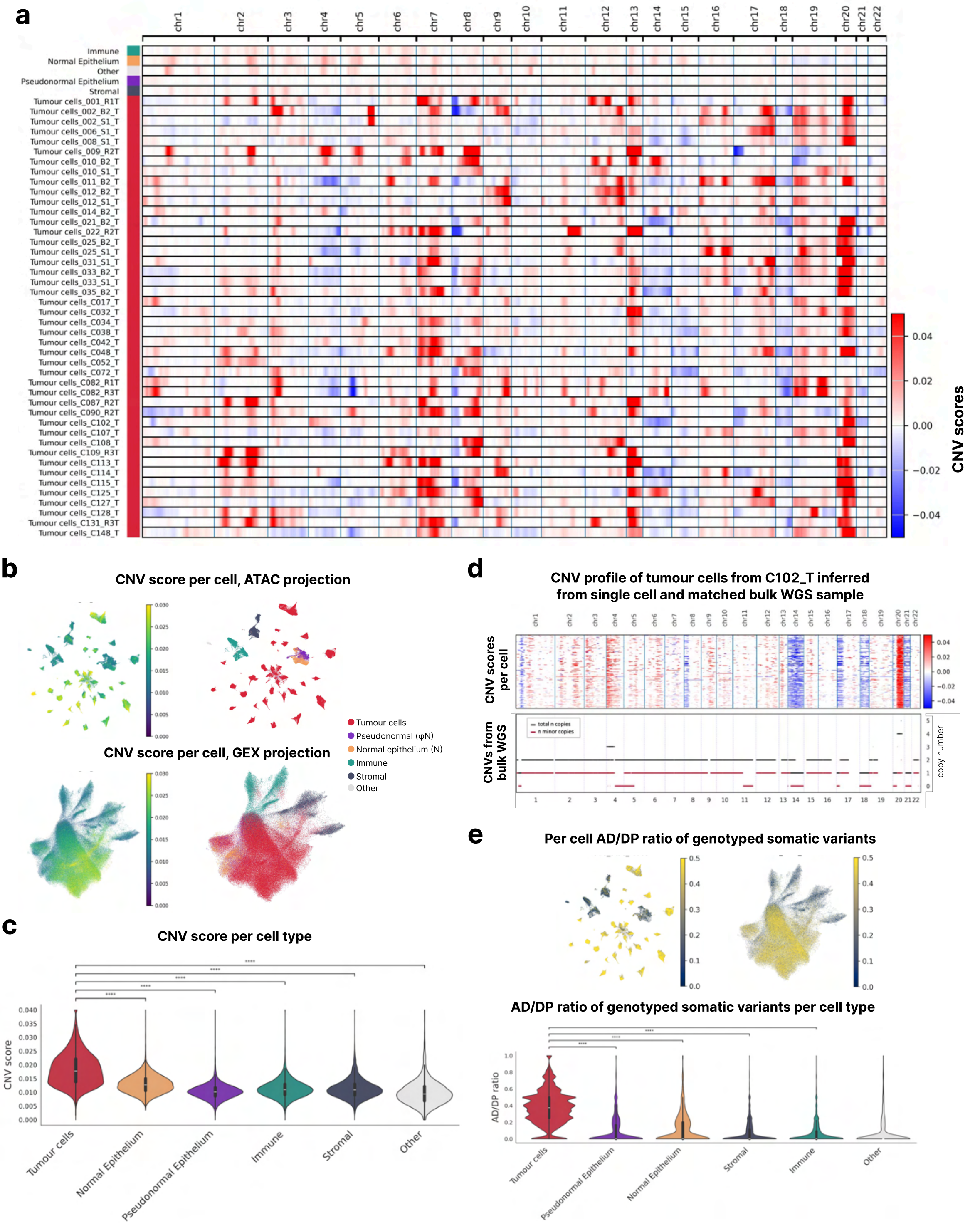


**Extended Data Figure 3. (a)** Copy number variations (CNVs) inferred from the single-cell cohort using InferCNVpy (GEX) with a 500-gene window, averaged per cell type. Blue indicates estimated copy-number loss, red indicates amplification related to diploid segment status. **(b)** UMAP projections of gene expression (GEX) and ATAC datasets, colored by per-cell CNV score (*left*) and cell type (*right*). **(c)** CNV burden per cell across cell types (per-cell CNV score calculated as the L2 norm of the cell CNV profile; see Methods). **(d)** Inferred CNVs in tumor cells from patient C102_T (*top*) compared with CNVs estimated by Sequenza from bulk whole-genome sequencing (*bottom*; see Methods). **(e)** Allelic depth (AD) / total depth (DP) ratios across genotyped somatic variants in scATAC-seq data, calculated per cell (*top*) and per cell type (*bottom*) using matched WGS samples (see Methods). Higher AD/DP ratios in tumor cells indicate elevated mutational burden.

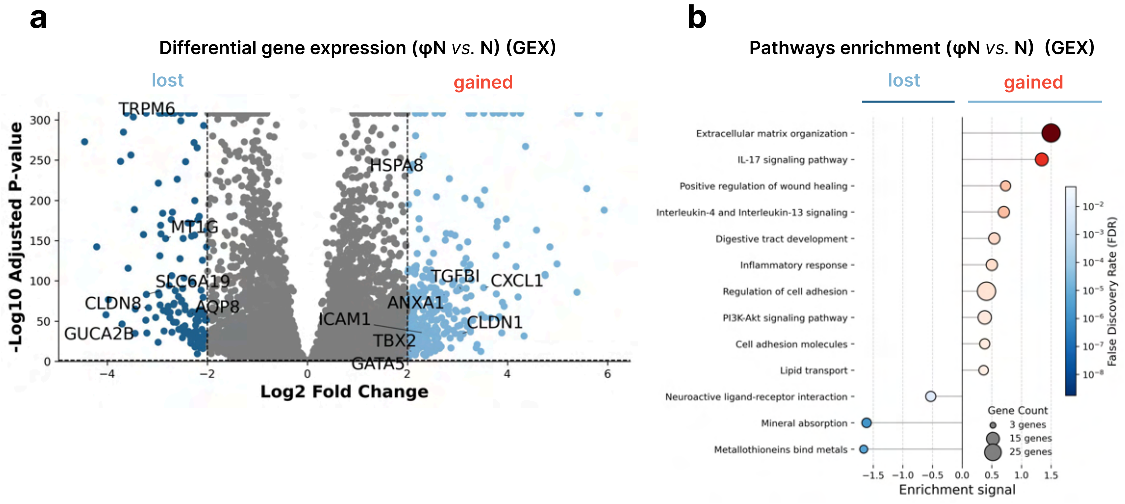


**Extended Data Figure 4. (a)** Genes differentially expressed in pseudonormal epithelial cells (light blue) vs. normal distant epithelial cells (deep blue). **(b)** STRING pathway enrichment analysis of genes upregulated in pseudonormal epithelial cells (red) vs. normal distant epithelial cells (blue).

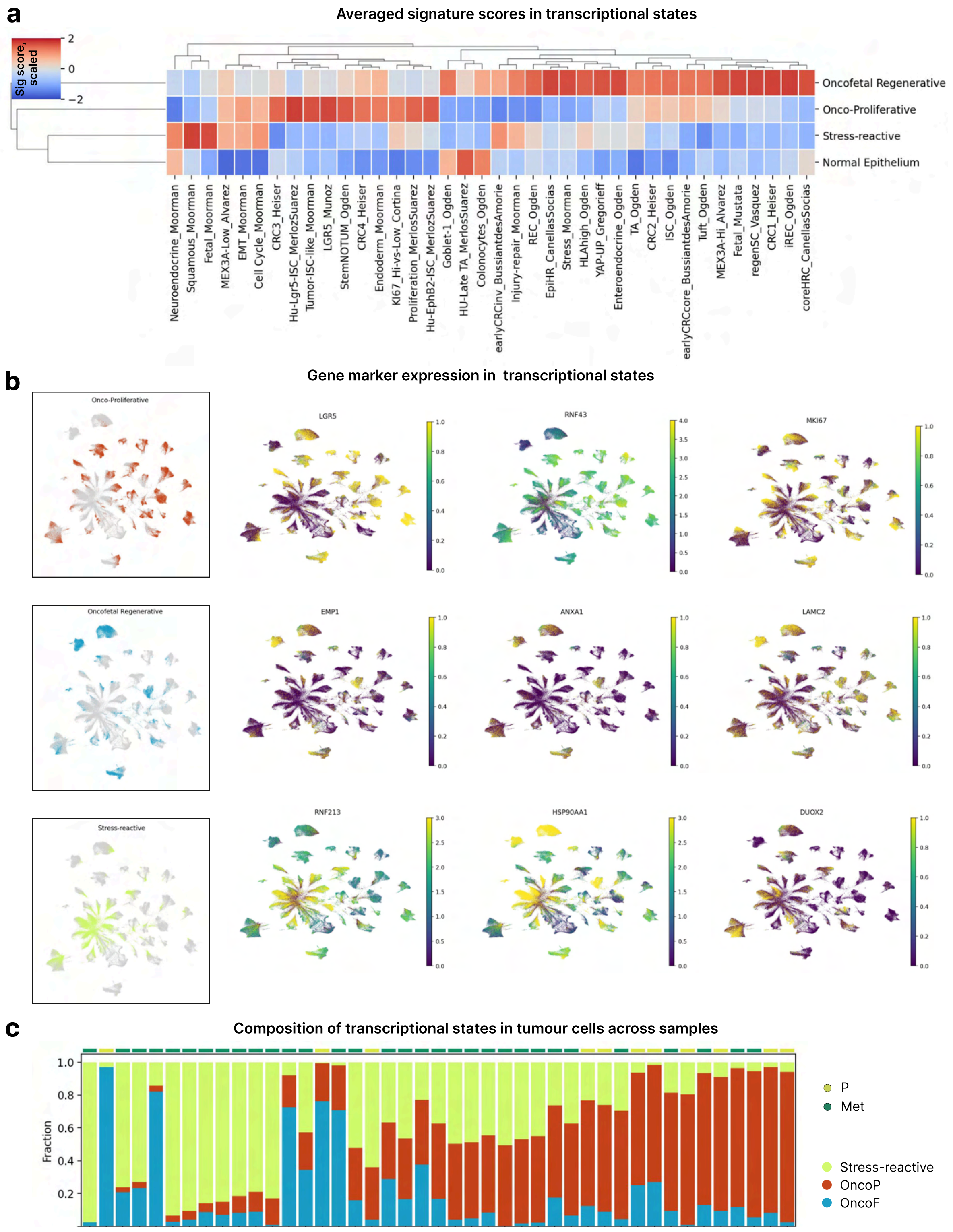


**Extended Data Figure 5. (a)** Mean scaled signature score for 38 reference signatures (see Extended Data Table X) used to cluster epithelial cells into transcriptional states. **(b)** UMAP visualization of epithelial cells showing the distribution of the three transcriptional states (*left* panels) and expression of representative marker genes (*right* panels) **(c)** Relative abundance of epithelial transcriptional states across individual samples (sorted by fraction of oncoproliferative state)

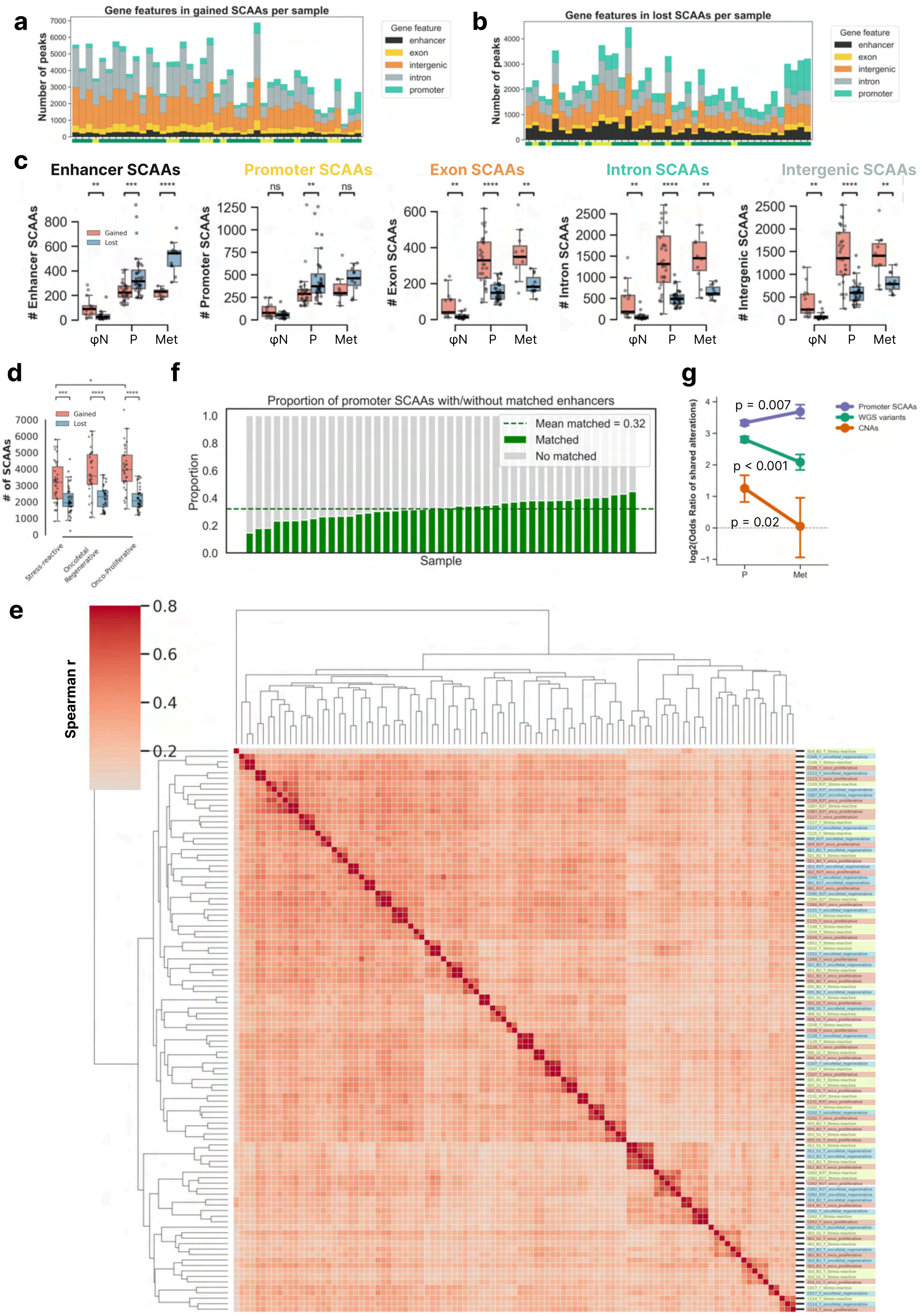


**Extended Data Figure 6.** **(a)** Genomic features associated with gained and **(b)** lost SCAAs in tumour cells compared with normal epithelial cells in primary and metastatic samples. **(c)** Number of gained (pink) and lost (blue) SCAAs associated with genomic features in pseudonormal epithelium, primary and metastasis CRC tumours **(d)** Number of gained and lost SCAAs (*left*) and total number of SCAAs per tissue type (*right*) across transcriptional states of epithelial cells **(e)** Pairwise similarity of tumor SCAA profiles across transcriptional states per sample. Transcriptional states are highlighted. Similarity is shown as Spearman correlation coefficients calculated using $\log_2$ fold-change values of SCAAs.**(f)** Fraction of promoter SCAAs with matched enhancer across samples **(g)** Averaged enrichment of shared genomic and epigenomic alterations between pairs of primary and metastatic tumours, measured as odds ratios for shared single-nucleotide variants, copy-number alterations and promoter-associated SCAAs. Points represent the mean $\log_2$ odds ratio across pairwise tumour comparisons, and error bars indicate 95% confidence intervals estimated by bootstrapping.

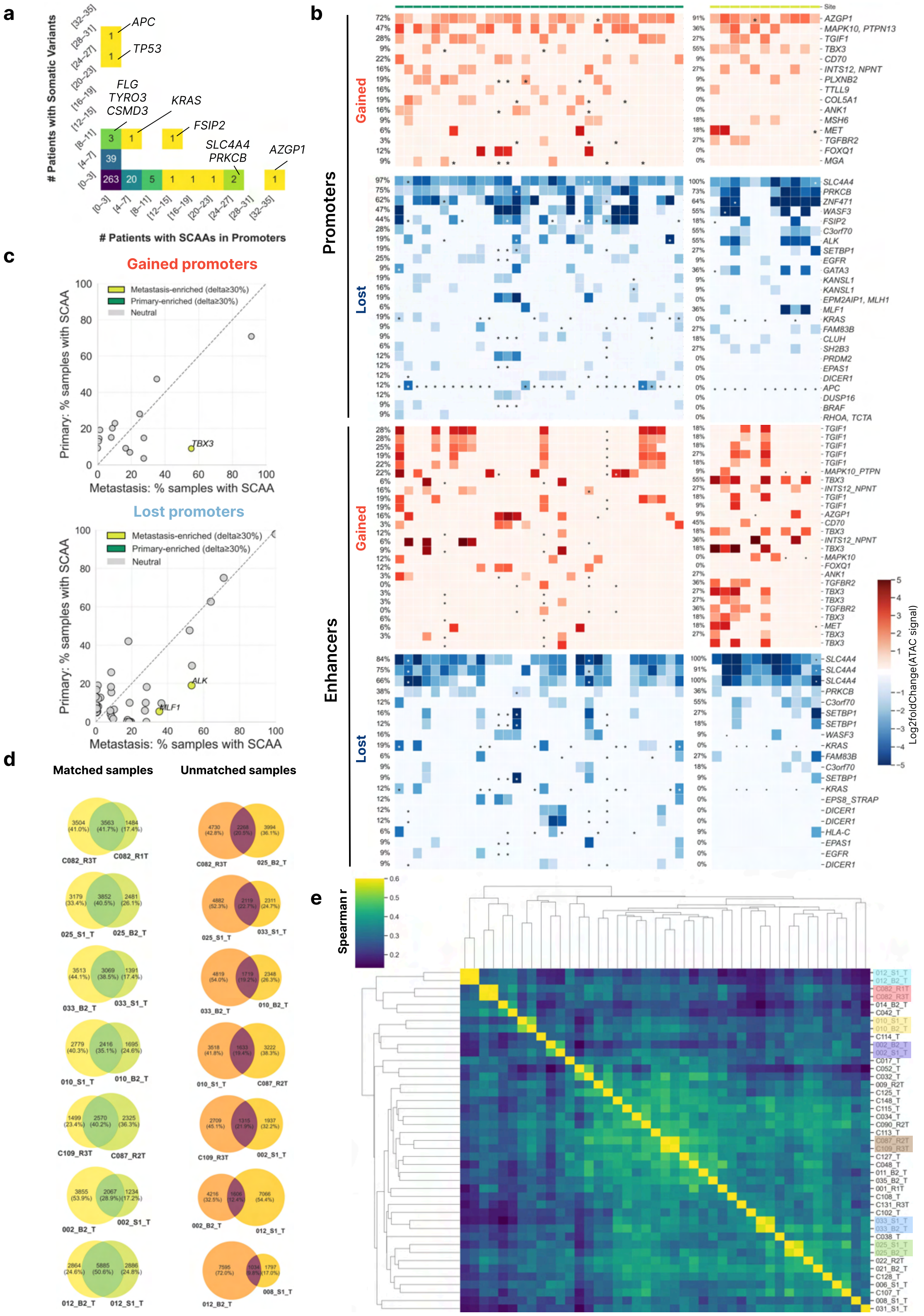

**Extended Data Figure 7. (a)** Relationship between genetic (somatic variants) and epigenetic (SCAAs) regulation across cancer driver genes. **(b)** Top recurrent SCAAs (|$\log_2$ fold change| > 1, padj < 0.01) affecting promoters and enhancers of cancer-related genes in CRC malignant cells relative to normal colon epithelial cells. Shown are SCAAs recurrent in at least three patients. Enhancers with a corresponding affected promoter are indicated. Asterisks denote the presence of somatic variants in the associated gene. Same heatmap for all genes is presented in Fig. 2b. **(c)** Recurrence of gained (*top*) and lost (*bottom*) promoter-associated SCAAs in primary tumors and metastatic samples. Genes enriched in metastatic SCAAs are annotated. **(d)** Number of shared and not shared SCAAs (|$\log_2$ fold-change| > 1, padj < 0.01) in seven matched (*left*) and seven random unmatched (*right*) sample pairs. **(e)** Pairwise similarity of tumor SCAA profiles across CRC samples. Matched samples from the same patient highlighted (see Extended Data Fig. 1 for sample annotation). Similarity is shown as Spearman correlation coefficients calculated using $\log_2$ fold-change values of SCAAs.

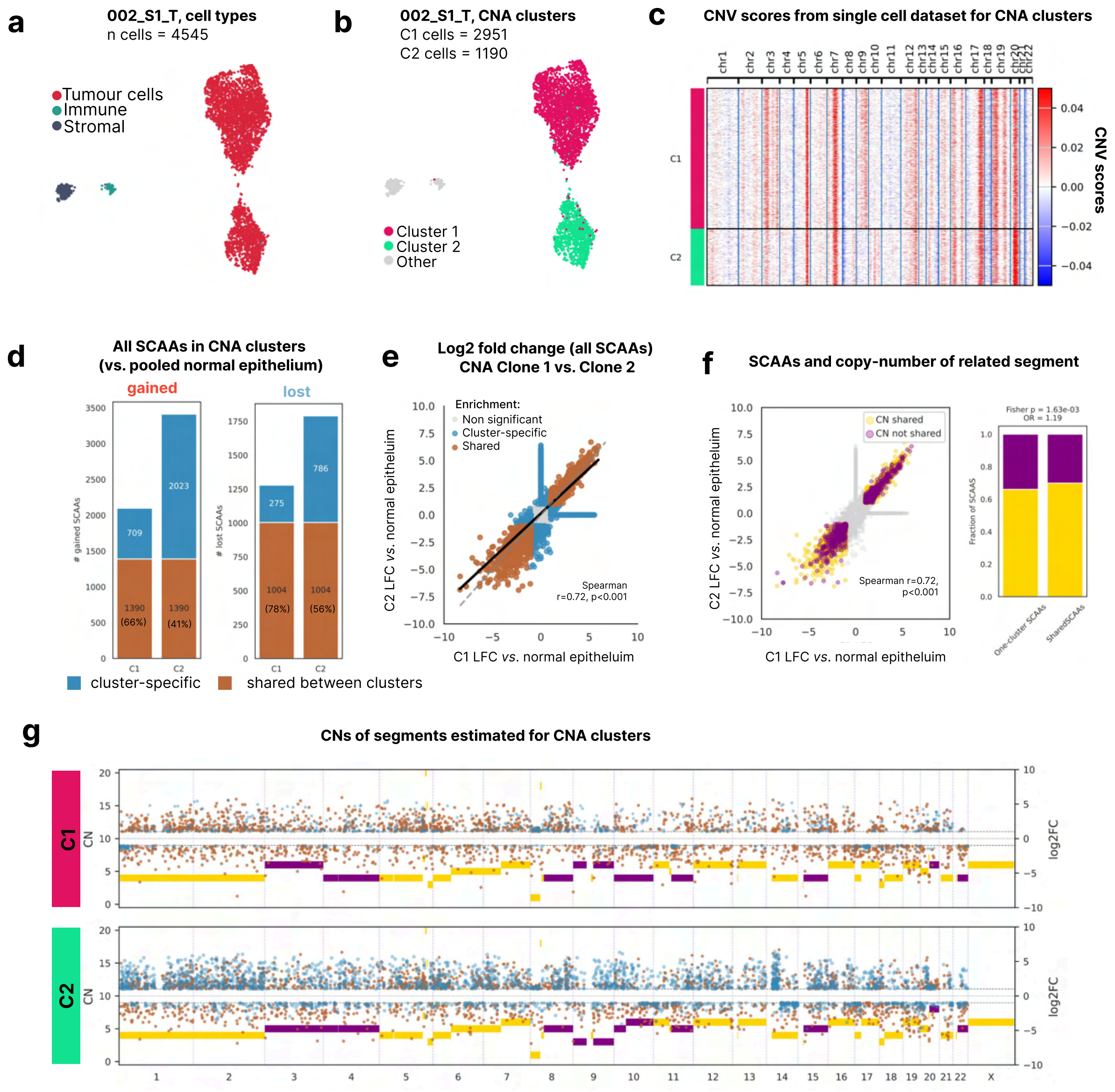


**Extended Data Figure 8. (a)** UMAP visualization of cells from sample 002_S1_T colored by cell type annotation. **(b)** UMAP visualization of cells from sample 002_S1_T with tumour cells colored by CNV clusters inferred using CONGAS+ (see Methods). **(c)** Single-cell copy number variation (CNV) scores of tumour clusters C1 and C2 inferred using InferCNVpy (GEX) with a 500-gene smoothing window. Blue indicates inferred copy-number loss and red indicates inferred copy-number gain relative to diploid reference segments. **(d)** Number of shared and cluster-specific SCAAs in C1 and C2 tumour clusters vs. pooled normal epithelial cells **(e)** Correlation of log2 fold-change values for differentially accessible peaks in C1 and C2 relative to normal epithelial cells. Shared and cluster-specific SCAAs are highlighted in brown and blue, respectively. SCAAs were defined as differentially accessible peaks with $|log2FC| > 1$ and adjusted $p < 0.01$. **(f)** *Left*, correlation of log2 fold-change values for differentially accessible peaks in C1 and C2 relative to normal epithelial cells. SCAAs mapping to genomic segments with shared copy-number states are highlighted in gold, whereas SCAAs associated with segments exhibiting different copy-number states between clusters are highlighted in purple. *Right*, fraction of shared and cluster-specific SCAAs associated with genomic segments displaying shared (gold) or differential (purple) copy-number states.**(g)** Copy number profile inferred in C1 and C2 clusters with CONGAS+ (see Methods). With dots are annotated coordinates of shared (brown) and cluster-specific (blue) SCAAs.

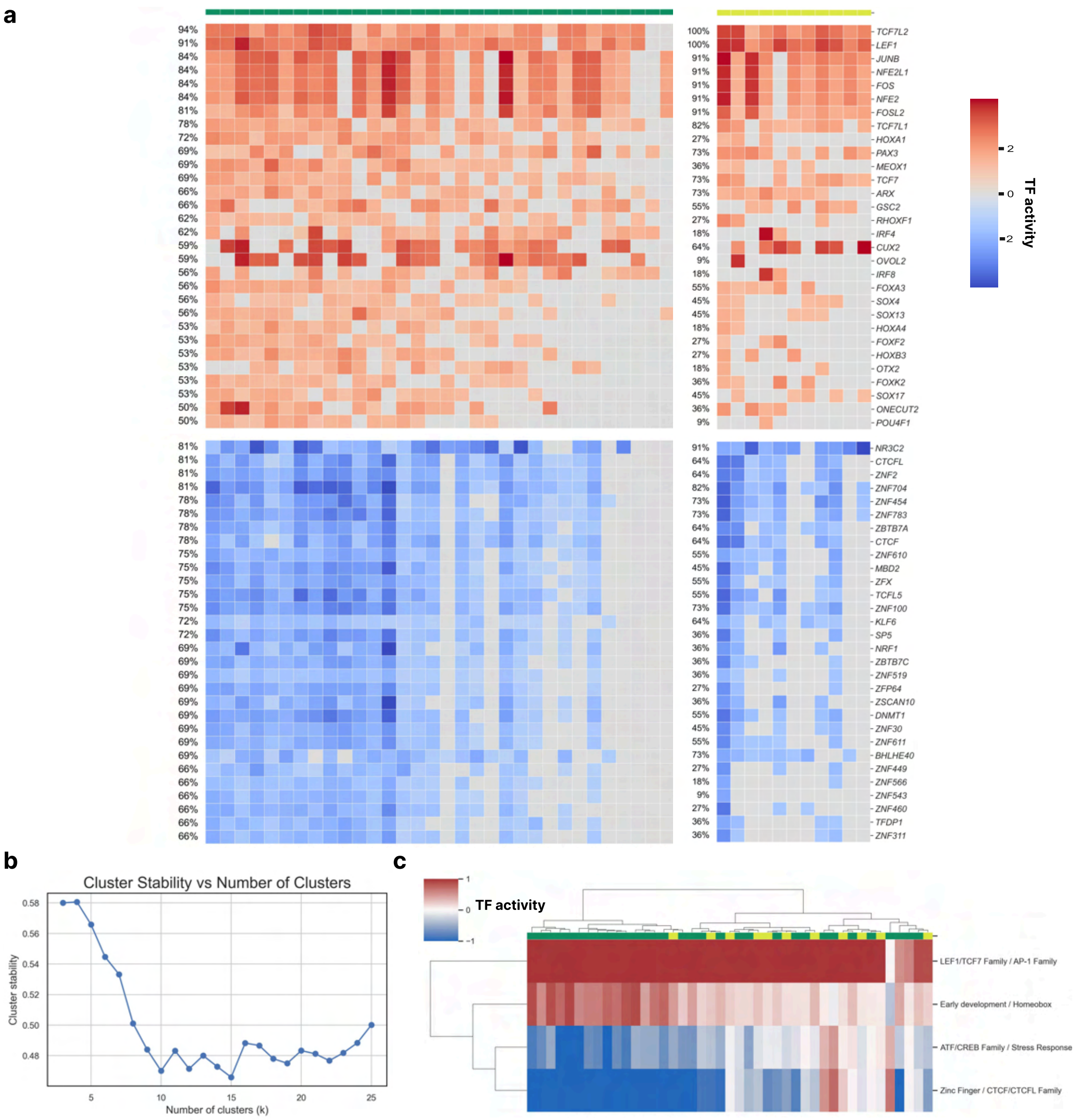

**Extended Data Figure 9. (a)** CIS-BP transcription factor motif enrichment analysis of transcription factor (TF) binding sites across gained and lost SCAAs. TFs that were significantly upregulated (red) or downregulated (blue) in at least nine patients are shown. **(b)** Stability scores assessing the robustness of TF clustering across different numbers of clusters ($k$ = 2–25). Stability was computed as the average pairwise consensus with which TF pairs co-clustered across 80 bootstrap iterations, each subsampling 80% of TFs (see Methods). Clustering was performed using hierarchical clustering with Ward linkage and Euclidean distance. **(c)** Per-sample averaged TF enrichment scores for each of the four identified TF clusters.

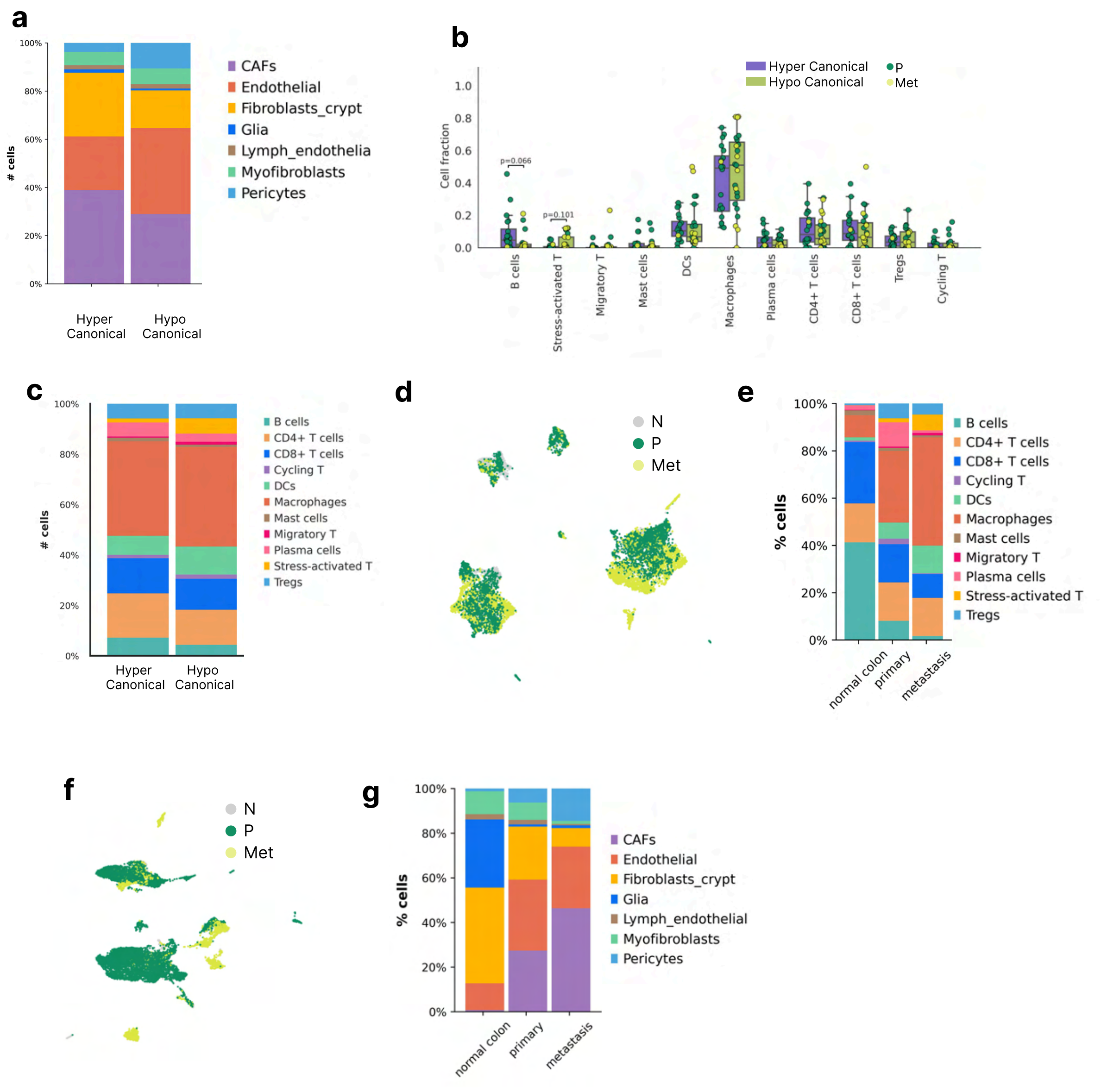


**Extended Data Figure 10. (a)** Distribution of stromal subpopulations across hyper- and hypocanonical groups of samples **(b)** Immune cells abundance in Hypo- and Hyper-canonical groups of tumour samples **(c)** Distribution of immune subpopulations across hyper- and hypocanonical groups of samples **(d)** UMAP projection of scATAC-seq data showing 12,021 immune cells across the CRC cohort; tissue site is highlighted **(e)** Distribution of immune subpopulations across normal colon, primary CRC, and metastatic CRC samples. **(f)** UMAP projection of scATAC-seq data showing 13,903 stromal cells across the CRC cohort; tissue site is highlighted **(g)** Distribution of stromal subpopulations across normal colon, primary CRC, and metastatic CRC samples.

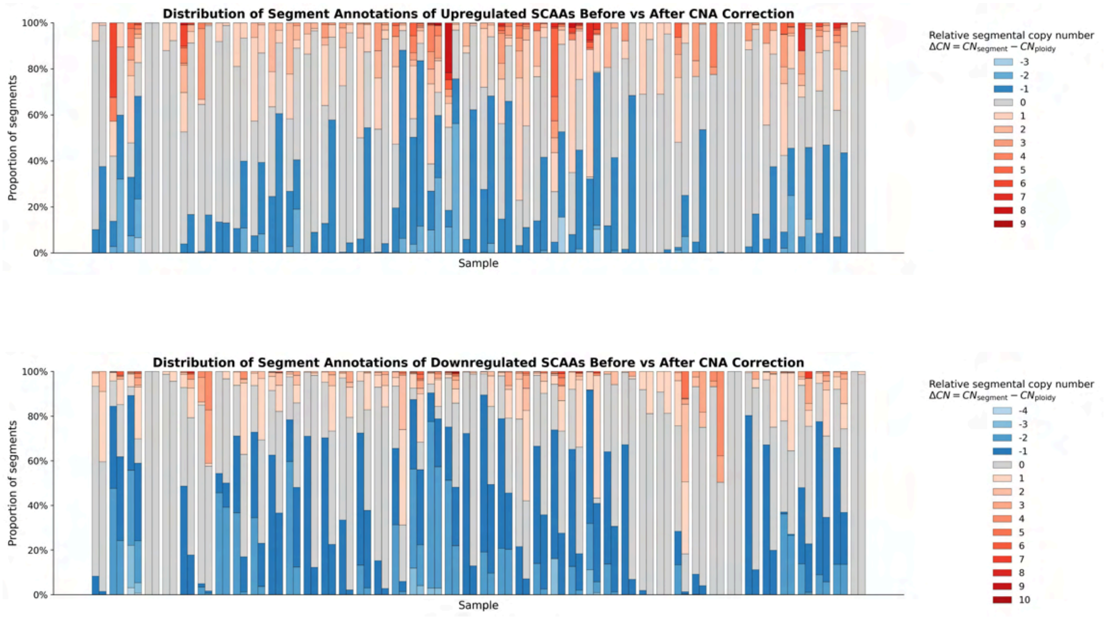


**Supplementary Figure 1. Fraction of SCAAs in every sample before (left bar) and after (right bar) correction of ATAC signal per copy-number**
